# Mitochondrial dysfunction activates a translational stress response that drives diabetic pain

**DOI:** 10.64898/2026.11.11.765941

**Authors:** Angela R. Meyer, Guadalupe García, Shea Ruhly, Alexander R. Mikesell, Jacob B. Shapiro, Rebecca Chase, Zachary T. Campbell

**Affiliations:** Department of Anesthesiology, University of Wisconsin – Madison, Madison, WI, 53705, United States; Department of Genetics, Washington University in St. Louis School of Medicine, St. Louis, MO, 63130, United States; Department of Psychiatry, Washington University in St. Louis School of Medicine, St. Louis, MO, 63130, United States

**Author notes:** Correspondence to: Zachary Campbell 600 Highland Avenue, Madison, WI, 53792.

**Keywords:** MGO, neuropathic pain, GCN2, ISR

## Abstract

Neuropathic pain is common and debilitating among diabetics. The reactive glycolytic metabolite methylglyoxal (MGO) is strongly associated with painful diabetic neuropathy. MGO triggers pain through activation of the integrated stress response (ISR). The underlying molecular mechanisms are unclear. Here, we show that general control nonderepressible 2 (GCN2) kinase acts as a conserved sensor that bridges glycolytic and mitochondrial dysfunction to pain hypersensitivity. In human sensory neurons derived from dorsal root ganglion (DRG), MGO increased eIF2α phosphorylation and suppressed global translation. These effects were blocked upon inhibition of GCN2. Unbiased ribosome profiling of mouse DRG revealed that MGO triggered repression of mRNAs that encode components of the electron transport chain. Based on this observation, mitochondrial dysfunction was examined as a potential upstream trigger for GCN2 activation. MGO directly glycated mitochondrial complex I and increased the NADH/NAD^+^ ratio. Correcting this redox imbalance via expression of a mitochondrial-targeted bacterial NADH oxidase (mitoLbNOX) prevented ISR activation by MGO. In a streptozotocin-induced mouse model of diabetic neuropathy, pharmacological GCN2 inhibition or sensory neuron-specific conditional deletion significantly attenuated mechanical allodynia. Calcium imaging revealed that genetic deletion of GCN2 reduced calcium influx in response to an agonist of TRPA1. Finally, expression of mitoLbNOX using adeno-associated viruses (AAV) in DRG sensory neurons *in vivo* reduced streptozotocin-induced mechanical hypersensitivity. Our findings demonstrate that GCN2 is a central, conserved driver of translational dysregulation downstream of mitochondrial redox stress in diabetic pain. This work highlights the critical role of mitochondrial function and translational control as key drivers of painful diabetic neuropathy.

## Introduction

Diabetes impacts 11% of the global population [77]. Over 100 million people with diabetes develop neuropathic pain [88]. Standard therapeutics for diabetic pain target the central nervous system (*e.g.,* gabapentinoids, SSRIs). A fundamentally different approach involves reversal of maladaptive changes in peripheral sensory neurons that are integral to the genesis and maintenance of chronic pain [64]. The major goal of this work is the identification of molecular mechanisms that drive diabetic pain.

A key metabolic alteration triggered by diabetes is the accumulation of the glycolytic metabolite MGO. MGO circulates at 100-250 nM in healthy individuals, but is increased to 0.5-5 µM in type I and type II diabetes [2; 8; 65]. MGO covalently reacts with proteins, lipids, and nucleic acids to form advanced glycation end products [10; 27; 51; 62; 63]. At pathological concentrations, MGO is sufficient to trigger pain through direct effects on sensory neurons, although much of the underlying mechanism is unclear [4; 6; 8; 24; 55].

A critical homeostatic translational control mechanism engaged by MGO is the ISR [6; 44; 55; 56; 90]. The ISR is initiated by four sensory kinases: GCN2, PERK, PKR, and HRI [85]. While each kinase is responsive to a distinct form of cellular stress, they converge on phosphorylation of eIF2α. Phosphorylation of eIF2α (p-eIF2α) inhibits the exchange of GDP for GTP by eIF2B and prevents delivery of the initiator Met-tRNA_i_-GTP ternary complex to ribosomes [74; 86]. The ISR has bifurcated effects on translation: most mRNAs are repressed, while transcripts containing upstream open reading frames (uORFs) are preferentially translated [33; 52; 82]. Chronic activation of the ISR promotes cell death while low-grade activation supports recovery [1; 61]. In neurons, the ISR influences excitability and plasticity [16; 17; 50; 76; 91].

GCN2 is the ancestral ISR kinase. It senses dysfunctional elongation in response to an array of cues that induce ribosome collisions (*e.g.,* amino acid depletion) [60; 68; 70; 72; 87]. Mitochondrial dysfunction has recently emerged as a GCN2 trigger [48; 57; 66]. In addition to ATP synthesis, the mitochondria facilitate metabolic recycling, which is key to amino acid catabolism. For example, complex I of the electron transport chain is a ubiquinone oxidoreductase that converts NADH to NAD^+^. Reduced NAD^+^ leads to depletion of the asparagine pool and causes GCN2 activation [57]. How metabolism intersects with translational control in the context of neuropathic pain is unknown.

Here, we identify GCN2 as responsible for activation of the ISR in human sensory neurons. Unbiased measurements of mRNA abundance and translation in DRG revealed widespread repression of ribosome biogenesis and electron transport chain components. Additionally, MGO directly modifies complex I of the electron transport chain resulting in accumulation of NADH. Metabolic reprogramming through the introduction of a bacterial NADH oxidase reversed the effects of MGO on the ISR and significantly diminished mechanical allodynia in diabetic mice. Finally, GCN2 is required for activation of the ISR and mechanical allodynia in a diabetic mouse model. We propose that GCN2 is a conserved bridge that links metabolic dysfunction to pain amplification.

## Methods

### Animals

Mice were housed in a 12-hour light/dark cycle with an ambient temperature of 20-26° C at 30-70% humidity. They were provided with access to food and water. To generate GCN2 conditional deletion animals, *GCN2*^fl/fl^ mice (JAX strain 008452) were crossed with *Pirt*^Cre/Cre^ mice [43; 53]. Wild-type 8–12-week-old C57BL/6 (B6) mice were obtained from the Biomedical Research Model Services at the School of Medicine and Public Health, University of Wisconsin-Madison. Male and female mice were tested separately to investigate sex differences. All animal procedures were approved by the Institutional Animal Care and Use Committee (protocol ID: M006582-A07) of the University of Wisconsin-Madison.

### Behavioral testing

Mechanical sensitivity was assessed using calibrated von Frey filaments (BIOSEB Instruments, BIO-VF-M, Pinellas Park, FL) and the up-down method as previously described [14; 23]. Animals were randomly assigned to treatment groups, and all behavioral testing was performed by an experimenter blinded to treatment conditions. All behavioral assessments were conducted during the light phase of the light-dark cycle between 0600 and 1800 hours.

Mice were placed individually in acrylic chambers positioned on an elevated wire mesh floor and allowed to acclimate for one hour prior to testing. Von Frey filaments were applied to the plantar surface of the hind paw for approximately two seconds, and paw withdrawal thresholds were calculated in grams (g) using the up-down method. Baseline mechanical sensitivity was assessed prior to AAV administration and again following intrathecal injection to ensure that AAV delivery did not alter mechanical sensitivity. Mice were allowed a two-week recovery and viral expression period prior to induction of diabetic pathology. Mechanical sensitivity was then assessed weekly following streptozotocin (STZ) administration on days 7, 14, 21, and 28.

### Induction of Diabetic Pathology

Two weeks following intrathecal AAV administration, diabetic pathology was induced using a multiple low-dose streptozotocin protocol [31]. Although female mice are known to be less sensitive to streptozotocin-induced β-cell toxicity than males, both male and female mice were included in this study to improve the generalizability of findings [45]. Thus, male and female mice were analyzed separately to account for potential sex-dependent differences in susceptibility to streptozotocin and the development of diabetic pathology.

Mice were fasted for four to six hours prior to each streptozotocin administration while maintaining free access to water. Streptozotocin (S0130-500MG, Sigma-Aldrich, St. Louis, MO) was freshly dissolved in 50 mM sodium citrate buffer (pH 4.5), protected from light, maintained on ice, and administered immediately following preparation. Mice received intraperitoneal injections of streptozotocin (50 mg/kg) once daily for five consecutive days. To reduce the risk of streptozotocin-induced hypoglycemia, mice were provided with 10% sucrose water following each injection throughout the treatment period and returned to regular drinking water one day after the final injection.

Body weight and blood glucose concentrations were monitored throughout the study to assess the development of diabetic pathology. Mice were considered diabetic and included in subsequent analyses if fasting blood glucose concentrations exceeded 200 mg/dL at four weeks following streptozotocin administration. This threshold was selected because mild hyperglycemia is characterized by fasting glucose concentrations greater than 150 mg/dL, and severe diabetes is often associated with blood glucose levels exceeding 300 mg/dL [28].

### Fed Blood Glucose Measurements

Blood glucose concentrations were measured to verify the development of streptozotocin-induced diabetes. Mice were fasted for four to six hours prior to sample collection with free access to water. Blood was collected from the tail vein and glucose concentrations were measured using an AlphaTrak 2 Veterinary Blood Glucose Monitoring System. Blood glucose measurements were obtained on days 7, 14, 21, and 28 following initiation of streptozotocin treatment, concurrent with behavioral testing and body weight measurements. Fasting blood glucose concentrations measured on day 28 were used to determine study inclusion, with mice required to exhibit glucose concentrations greater than 200 mg/dL to meet criteria for diabetic pathology.

### AAV Injections

rAAV encoding hSyn::DIO::mitoLbNOX::P2A::eGFP or hSyn::DIO::eGFP were delivered by intrathecal injection. 3.5 x 10^11^ viral genomes were administered in a total volume of 10 μL of sterile-filtered Takara Suspension Buffer. Mice were anesthetized with 2-3% isoflurane in oxygen delivered through a nose cone. The L4-L5 intervertebral space was identified anatomically, and a Hamilton syringe fitted with a 31-gauge needle was inserted into the intrathecal space. Successful intrathecal delivery was confirmed by observation of a characteristic tail-flick reflex. Following injection, mice were monitored until fully recovered from anesthesia and returned to their home cages. Mice were allowed a two-week recovery period to permit trans gene expression prior to induction of diabetic pathology with streptozotocin.

### Primary DRG cultures

Primary mouse sensory neurons were cultured as previously described [70; 71]. Briefly, 8-12-week-old Swiss Webster mice were deeply anesthetized with 5% isoflurane and euthanized by decapitation. The vertebral column was dissected and hemisected, the spinal cord and dura were removed, and the DRGs were collected in cold HBSS. DRGs were pelleted by centrifugation (400 × g, 2 minutes), the HBSS was aspirated, and the tissue was resuspended in 1 mg/mL collagenase A (Roche, Basel, Switzerland) in HBSS. DRGs incubated for 25 minutes at 37° C. The tissue was pelleted again (400 × g, 30 seconds) and the collagenase A was removed. DRGs were then resuspended in HBSS containing 1 mg/mL collagenase D (Roche) and 10% (v/v) papain (Roche) and incubated for 20 minutes at 37° C. After another centrifugation (400 × g, 30 seconds), the tissue was resuspended in DMEM/F12+GlutaMAX (Gibco) containing 1 mg/mL trypsin inhibitor (Roche) and 1 mg/mL BSA, and mixed until homogeneous. The suspension was passed through a 70 µm cell strainer into a 50 mL conical tube. The strainer was rinsed dropwise with 15-20 mL of DMEM/F12+GlutaMAX. The filtrate was centrifuged (400 × g, 5 minutes), the medium was aspirated, and the cells were resuspended in an appropriate volume of DRG medium (DMEM/F12+GlutaMAX, 10% FBS, 1% penicillin/streptomycin, and 2 ng/mL 5-fluoro-2’-deoxyuridine).

For immunofluorescence, cells were plated on 8-well chamber slides and cultured for six days (LabTek, Scotts Valley, CA, USA). For calcium imaging, cells were plated on 12 mm German Glass Coverslips (Fisher Scientific, Waltham, MA) and cultured for 24-48 hours. For both applications, slides and coverslips were coated with 1 µg/cm² poly-D-lysine (Gibco) diluted in PBS for 2 hours at 37° C prior to plating.

### HD10.6 cultures

HD10.6 cells were obtained from Ian Mohr (Department of Microbiology, NYU). Cells were grown in complete proliferating base media (PBM). PBM consists of advanced DMEM/F12 (12634010, Gibco, Waltham, MA) supplemented with Glutamax (35050061, ThermoFisher Scientific, Waltham, MA). Complete PBM is composed of 50 mL PBM, 1 mL Neurocult SM1 (100-1281, Stem Cell Technologies, Vancouver, Canada), 50 µL Prostaglandin E1 (P5515-1MG, Sigma-Aldrich), 100 µL Primocin (ant-pm-05, Invivogen, San Diego, CA), and 5 ng/mL β-FGF (100-18B, Stemfactor, Beltsville, MD). Prior to plating, 6-well plates were coated with 17 µg/mL fibronectin (FC010-5MG, Sigma-Aldrich) for one hour. TrypLE (12605010, ThermoFisher Scientific) was used to trypsinize cells. To differentiate HD10.6 cells, cultures were plated on 8-well chamber slides. Prior to plating, chamber slides were coated with 50 µg/mL poly-L-ornithine (Sigma, P3655-10MG) diluted in 0.5 M borate buffer (J60068, ThermoFisher Scientific) overnight. The following day, slides were washed three times with sterile water and slides were coated with 1 µg/mL fibronectin. Before plating, slides were washed with sterile water once and allowed to fully air dry. HD10.6 cells were plated at 30% confluency in complete PBM. The following day, the media was exchanged with differentiation media. Base media consisted of Neurobasal (21103049, Gibco) supplemented with Glutamax. Complete differentiation media consisted of 50 mL of base media, 1 mL B27 supplement (17504001, ThermoFisher Scientific), 1 µg/mL doxycycline (T7660, Sigma-Aldrich), 50 ng/mL β-NGF (A42578, ThermoFisher Scientific), 25 ng/mL CNTF (450-13-100ug, ThermoFisher Scientific), 25 ng/mL GDNF (450-10-10UG, ThermoFisher Scientific), 25 µM forskolin (11018-10, Cayman chemical, Ann Arbor, MI), and 100 µL Primocin. Cells were differentiated for at least seven days prior to experiments.

### F11 and HEK293T cell culture

F11 cells (ECACC 8062601, RRID:CVCL_H605) and HEK293T cells were maintained at 37° C in a humidified incubator with 5% CO2. Media consisted of DMEM (Gibco, 11-995-073) with 10% fetal bovine serum (FBS) as described [70]. Cells were passaged every 2-4 days as needed. All cell culture procedures were performed under sterile conditions in a biosafety cabinet. For differentiation, F11 cells were cultured in DMEM + 1% FBS with 30 µM forskolin for 72 hours. For transfections, cells were grown to 70% confluency in 8-well chamber slides. For one well, 0.5 µL Lipofectamine 3000 was diluted in 15 µL Opti-MEM (31985062, ThermoFisher Scientific). In a separate tube, 0.5 µL P3000 and 250 ng mitoLbNOX were diluted in 15 µL Opti-MEM. The two tubes were mixed and incubated at room temperature for 15 minutes. The total 30 µL was added to each well. The following day, the media was replaced with differentiation media.

### Ribosome profiling

MGO (M0252, Sigma-Aldrich) was injected into mice intraperitoneally (2 mg/kg) for six days as described [6]. A separate cohort of animals was treated with saline and used as controls. Three hours after the last dose of MGO, dorsal root ganglia (DRG) from cervical (C1) to lumbar (L5) spinal segments were rapidly harvested and placed in ice-cold 1x HBSS (24-020-117, Gibco) supplemented with 2 mg/mL emetine. DRGs from 4 mice were pooled into 1 biological replicate. There were 4 biological replicates per treatment group. Following dissection, DRGs were transferred to ice-cold polysome lysis buffer containing 20 mM Tris-HCl, pH 7.5, 250 mM NaCl, 15 mM MgCl_2_, 1 mM DTT, 0.5% (vol/vol) Triton X-100, 2.5 U/mL DNase I, 40 U/mL RNasin and 50 µg/mL emetine, and then homogenized using a pre-chilled Dounce homogenizer.

Libraries consisting of ribosome-bound RNA fragments were generated as previously described with minor modifications [35; 36]. MicroSpin S-400 columns (GE Healthcare) were used to isolate ribosome-bound RNAs. After rRNA was removed using the RiboCop rRNA depletion kit (Lexogen), footprints were dephosphorylated and then size selected (28-34 nt) by PAGE on 15% TBE-Urea gels (Bio-Rad). Footprints were generated by SMARTer smRNA-Seq kit for Illumina (TaKaRa). RNA abundance was quantified using the QuantSeq 3ʹ mRNA-Seq library kit (Lexogen). The concentrations of purified libraries were quantified using Qubit (Invitrogen) and the average size was determined by fragment analyzer with high sensitivity NGS fragment analysis kit (Advanced Analytical Technologies Inc.). Libraries were then sequenced on an Illumina NextSeq500 sequencer using 75-bp single-end high output reagents (Illumina).

Data were subjected to a quality check using FastQC 0.11.5 (Babraham Bioinformatics). Adapters were subjected to trimming and mapping was conducted with TopHat 2.1.1 (with Bowtie 2.2.9) to the mouse reference genome (NCBI reference assembly GRCm38.p4) and reference transcriptome (GENCODE vM10). Strand orientation was considered during the mapping process and processed BAM files were quantified for each gene using Cufflinks 2.2.1 with GENCODE vM10 genome annotation. Read counts were not normalized by length using the Cufflinks option -- no–length–correction. Relative abundance for the i^th^ gene was determined by calculating TPM (transcripts per million) values as follows:

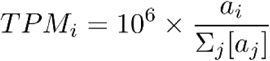

where a_j_ is the Cufflinks reported relative abundance.

Finally, TPM values were normalized to upper decile for each biological replicate and udTPM values (upper decile TPM) were used for analysis, to provide uniform processing for samples with varying sequencing depth [30].

### Immunofluorescence

Following treatments, cells were washed with ice-cold 1x phosphate-buffered saline (PBS). PBS was aspirated and 4% formaldehyde in PBS was added for fifteen minutes. Slides were washed with three two-minute PBS washes. To permeabilize, 0.5% Triton X-100 was added to cells for ten minutes. Then, cells were washed three times with two-minute PBS washes. Next, cells were blocked in 10% goat serum (G9023-10ML, Sigma-Aldrich) for one hour. Then, primary antibodies were added and cells were incubated overnight at 4° C on a rocker. The following primary antibodies were used: anti-peripherin (NBP1-05423, 1:1000, Novus, Centennial, CO) and anti-p-eIF2α (3398S, 1:200, CST, Danvers, MA). The next day, primary antibody was removed and three two-minute PBS washes were performed. Secondary antibody was added and cells were incubated for one hour at room temperature protected from light. The following secondary antibodies were used: anti-chicken Alexa Fluor 488 (A11039, 1:1000, Invitrogen) and anti-rabbit Alexa Fluor 568 (A11011, 1:1000, Invitrogen). Secondary antibody was removed and three two-minute PBS washes were performed. Coverslips were mounted onto the slides with Prolong glass anti-fade mounting media (P36984, ThermoFisher Scientific). Slides were sealed with clear nail polish.

### Puromycylation

For puromycylation, cells were treated with 1 µM puromycin (35841-10, Cayman chemical) for five minutes. For nucleolar puromycin staining, cells were treated with 1 µM puromycin for ten minutes. Following puromycin treatment, cells were washed with cold PBS + 0.00036% digitonin. Fixation and the following steps were performed as described in the immunofluorescence section above. The primary antibodies used were anti-peripherin (NBP1-05423, 1:1000, Novus), anti-puromycin (MABE343, 1:5000, EMD Millipore, Burlington, MA), and anti-nucleolin (14574T, 1:1500, CST). Anti-chicken Alexa Fluor 488 (A11039, 1:1000, Invitrogen, Waltham, MA), anti-mouse Cy5 (A10524, 1:1000, Invitrogen), and anti-rabbit 568 (A11011, 1:1000, Invitrogen) were the secondary antibodies used. For nucleolar puromycin staining, cells were incubated in DAPI for 10 minutes prior to mounting.

### Proximity ligation assay (PLA)

Cells were fixed and permeabilized as described above. Next, samples were treated with Duolink blocking buffer for one hour at 37° C. Cells were incubated in primary antibodies and rocked overnight at 4° C. The next day, PLA steps were performed according to the manufacturer’s instructions with minor adjustments (DUO92013, Sigma-Aldrich); wash buffer A was replaced with Tris-Buffered Saline with 0.1% Tween-20 (TBST) and wash buffer B was replaced with TBS. The following primary antibodies were used: anti-peripherin (Novus, 1:1000, NBP1-05423), anti-AGE (Abcam, 1:100, ab23722) and anti-complex I (Invitrogen, 1:50, 43-8800). Secondary probes were rabbit (DUO92002, Sigma-Aldrich) and mouse (DUO92004, Sigma-Aldrich). Anti-chicken Alexa Fluor 488 (A11039, 1:1000, Invitrogen) secondary antibody was used to detect peripherin.

### Immunohistochemistry (IHC)

DRG tissues were harvested following anesthetization with 2% isoflurane and transcardially perfused with ice-cold PBS, followed by 4% formaldehyde (in PBS, pH = 7.4). Tissues were harvested and postfixed overnight at 4° C with 4% formaldehyde. Then, tissues were cryoprotected in 30% sucrose for three days at 4° C. Tissues were embedded in Tissue-Tek OCT compound (Sakura Finetek, Torrance, CA) and cryosectioned at 14 µm. Tissue sections were washed to remove OCT compound and then incubated with 10 mM sodium citrate buffer for one hour. Tissues were then blocked with 10% normal goat serum with 0.3% Triton X-100 for two hours at room temperature using a humidified chamber. For GCN2 validation, goat anti-mouse Fab fragment (115-001-003, 1:1000, Jackson ImmunoResearch, West Grove, PA) was added to blocking solution. Sections were incubated overnight at 4° C with the following antibodies diluted in 1% normal goat serum in TBS plus 0.3% Triton X-100. For p-eIF2α analysis, anti-p-eIF2α (3398S, 1:150, CST), IB4 (I21412, 1:300, Invitrogen), and anti-TRPV1 (PA1-29770, 1:200, Invitrogen) were used. For GCN2 validation, anti-GCN2 (sc-374609, 1:200, Santa Cruz Biotechnology, Dallas, TX) and anti-peripherin (NBP1-05423, 1:1000, Novus) were used. Sections were washed three times with 0.1% Triton X-100 in TBS and incubated with the following secondary antibodies: anti-guinea pig 488 (A11073, 1:1000, Invitrogen) and anti-rabbit Cy5 (A10523, 1:1000, Invitrogen) for p-eIF2α analysis or anti-chicken Alexa Fluor 488 (A11039, 1:1000, Invitrogen) and anti-mouse Cy5 (A10524, 1:1000, Invitrogen) for GCN2 validation. Secondary antibodies were diluted in antibody solution for one hour at room temperature. After three washes, coverslips were mounted on the slides using ProLong Glass antifade.

### Imaging and quantification

Images were collected using a Leica SP8 laser scanning confocal microscope and a Nikon Eclipse TE200 inverted microscope. Z-stack images were captured using a 20X objective on the Leica and a 40X objective on the Nikon. Image quantification was performed using Fiji. A region of interest (ROI) was drawn around the peripherin-positive cells. Area, mean, and integrated density were measured and corrected total cell fluorescence (CTCF) was calculated with the following formula:

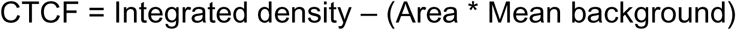

For PLA quantification, the number of puncta per cell was counted and puncta density was calculated as the number of puncta / area of the peripherin-positive cell. CTCF and PLA measurements were normalized to the vehicle average.

Images from immunohistochemistry experiments were captured on a Leica SP8 laser scanning confocal microscope with LAS X Life Science software. Images were acquired with a 20X objective, a pinhole of 1 Airy unit, and a scan speed of 400 Hz, using sequential acquisition. Representative images are presented as projections of Z stacks. To determine the p-eIF2α CTCF, ROIs were drawn around IB4^+^ or TRPV1^+^ neurons. The area, mean gray value, and integrated density were measured for each ROI. Corrected total cell fluorescence (CTCF) from the p-eIF2α channel was calculated and scaled to the vehicle group. For validation of the *GCN2*^fl/fl^ *Pirt*^Cre/+^ line, the number of cells that were GCN2 positive was divided by the total number of peripherin-positive neurons.

### Calcium Imaging

Calcium imaging experiments were carried out on DRG neurons 24-48 hours post-plating. Cells were loaded with the ratiometric calcium indicator Fura-2-AM (2.5 µg/mL) in 2% BSA for 45 minutes at room temperature. After loading, neurons were washed with an extracellular recording solution (pH 7.4) composed of 150 mM NaCl, 10 mM HEPES, 8 mM glucose, 5.6 mM KCl, 2 mM CaCl₂, and 1 mM MgCl₂. Imaging was performed on a Nikon Eclipse TE200 inverted microscope with image acquisition controlled by Nikon Elements software (Nikon Instruments, Melville, NY, USA). A RETRA Light Engine (Lumencor) provided excitation at 340 nm and 380 nm. Neuronal responsiveness to TRPA1 was evaluated by applying 10 µM ASP7663 (HY-101907, Medchem Express, Monmouth Junction, NJ) in the extracellular buffer, delivered for 120 seconds via gravity-fed perfusion at a rate of 1 mL per 10 seconds. Near the end of each recording, a 50 mM KCl stimulus served as a positive control to verify cell viability and responsiveness. Neurons were classified as responsive and were used for analysis only if their 340/380 fluorescence ratio increased by at least 20% in response to 50 mM KCl. Neurons failing this criterion were excluded from analysis. Neurons with a negative response to ASP7663 (less than zero) were excluded from analysis.

### NADH/NAD^+^ assay

HD10.6 cells were plated on 96-well white-bottom plates. After treatments, the Promega NAD/NADH-Glo assay was used according to the manufacturer’s instructions (G9072, Promega). Luminescence was detected using a Synergy H1 plate reader. NADH/NAD ratio was calculated as NADH luminescence / NAD luminescence.

### ATP/ADP assay

HD10.6 cells were plated on 96-well white-bottom plates. After treatments, the Sigma ATP/ADP ratio assay kit was used according to the manufacturer’s instructions (MAK135-1KT, Sigma-Aldrich). Luminescence was detected using a Synergy H1 plate reader. ATP/ADP ratio was calculated as ATP luminescence / ADP luminescence.

### LbNOX cloning and AAV preparation

MitoLbNOX was PCR amplified from Addgene #74448. MitoLbNOX was cloned into a piggybac plasmid backbone (Addgene #187950) via Gibson assembly to generate piggybac-mitoLbNOX-P2A-BFP for F11 transfection experiments. For AAV experiments, mitoLbNOX and P2A sequences were cloned into pAAV-hSyn-DIO-EGFP plasmid backbone (Addgene #50457) by Gibson assembly. Clones were confirmed by sequencing.

### rAAV Production and Delivery

70-80% confluent HEK293T cells were transfected with 3.75:1 PEI (linear, MW 25,000):DNA ratio. Immediately before transfection, media was changed to 5% FBS, 1x Non-Essential Amino Acids (11140050, Gibco). 5.7 μg rAAV genome plasmid, 11.4 μg pHelper (V005569, Novapro), and 22.9 μg pUCmini-MaCPNS1 (Addgene 185136) (40 μg total, 1:4:2 ratio) were delivered to each 15 centimeter plate. Media changes were performed 16 hours and 64 hours after transfection. rAAV particles were collected using the Takara AAVpro Purification Kit Maxi (6666) following the manufacturer’s protocol.

Viral genome copy number was quantified by RT-qPCR using two primer sets targeting distinct elements of the rAAV genome plasmid. Viral genome copy number was quantified as the average copy number from both assays. Primers used were WPRE (forward 5′-GGCTGTTGGGCACTGACAAT-3′, reverse 5′-CCGAAGGGACGTAGCAGAAG-3′), hGH (forward 5′-GGTGCGAGAAATGCCTAGAAGG-3′, reverse 5′-TGTCTTCCCAACTTGCCCCTT-3′). Linearized rAAV genome plasmid was used as a standard for copy-number quantification. DNase I was used to digest unencapsulated DNA. 2 µL of rAAV samples were diluted in 100 µL of 0.05 U/mL DNase I (EN0521, Thermo Scientific), 2 mM CaCl_2_, 10 mM Tris-HCl, 10 mM MgCl_2_ and digestion was carried out at 37° C for one hour. DNase I was inactivated by the addition of 25 mM EDTA and 70° C incubation for ten minutes. To liberate encapsulated DNA, 120 µL of 100 µg/mL Proteinase K (03115828001, Roche), 1 M NaCl, 1% wt/vol N-lauroylsarcosine sodium salt was added to samples. Digestion was carried out at 50° C for two hours. Proteinase K was inactivated by incubation at 95° C for ten minutes. Samples were diluted 1:300 in ultra-pure water prior to preparing the qPCR reaction with iTaq Universal SYBR Green Supermix (BioRad 1725121). qPCR was performed on an Applied Biosystems Real-time PCR StepOne Plus machine with the following program: initial denaturation at 95° C for ten minutes, followed by 40 cycles of 95° C for 20 seconds (denaturation) and 60° C for one minute (annealing/extension), with fluorescence reads after every cycle. Melt curve analysis was performed to assess amplicon specificity.

### Statistical analyses

The data in this study are reported as the mean ± standard error of the mean (SEM). For cell-based experiments, a One-way Analysis of Variance (ANOVA) followed by Sidak’s or Dunnett’s post hoc tests were applied. For all behavioral analyses, a Two-way ANOVA followed by Bonferroni’s post hoc test for multiple comparisons was applied. A statistical threshold of P < 0.05 was used to determine significance. All statistical analyses were conducted using GraphPad Prism v10.6.1.

## Results

### GCN2 activates the ISR in human neurons

Our prior work identified GCN2 as responsible for MGO-mediated induction of the ISR in primary murine DRG neurons [55]. To examine if the role of GCN2 in cellular responses to MGO is conserved, immortalized human embryonic dorsal root ganglion clonal cell line HD10.6 cells were differentiated and treated with MGO for 24 hours and p-eIF2α immunofluorescence was measured. MGO induced a modest but significant increase in p-eIF2α as compared to a vehicle group (**Fig. 1A**) (One-way ANOVA, F_3,442_ = 14.14, Sidak’s multiple comparison test, \**P* < 0.05). This finding is consistent with our prior observations in murine sensory neurons [55]. Blockade of GCN2 with a highly selective inhibitor (IN-7) [39] prevented activation of the ISR by MGO (**Fig. 1A**). We also tested whether blockade of PERK rescues ISR activation [6]. However, blockade of PERK did not suppress ISR activation in differentiated HD10.6 cells (**Fig. 1A**). Next, we examined the effects of MGO on global translation. Given that phosphorylation of eIF2α results in a global suppression of protein synthesis, we hypothesized that GCN2 was required for translational suppression by MGO [61]. Our approach makes use of the small molecule puromycin, which mimics the 3’ end of an aminoacylated tRNA and is readily incorporated into nascently translating polypeptides [67]. We found that MGO suppressed translation by 24% (**Fig. 1B**) (One-way ANOVA, F_3,480_ = 29.20, Sidak’s multiple comparison test, \*\*\*\**P* < 0.0001, n.s. = not significant). Blockade of GCN2 with IN-7 prevented translational repression by MGO, while inhibition of PERK had no effect (**Fig. 1B**). We conclude that repression of protein synthesis by MGO is GCN2-dependent. Collectively, these observations indicate that MGO induces the ISR via GCN2 in human sensory neurons.

**Fig. 1.**
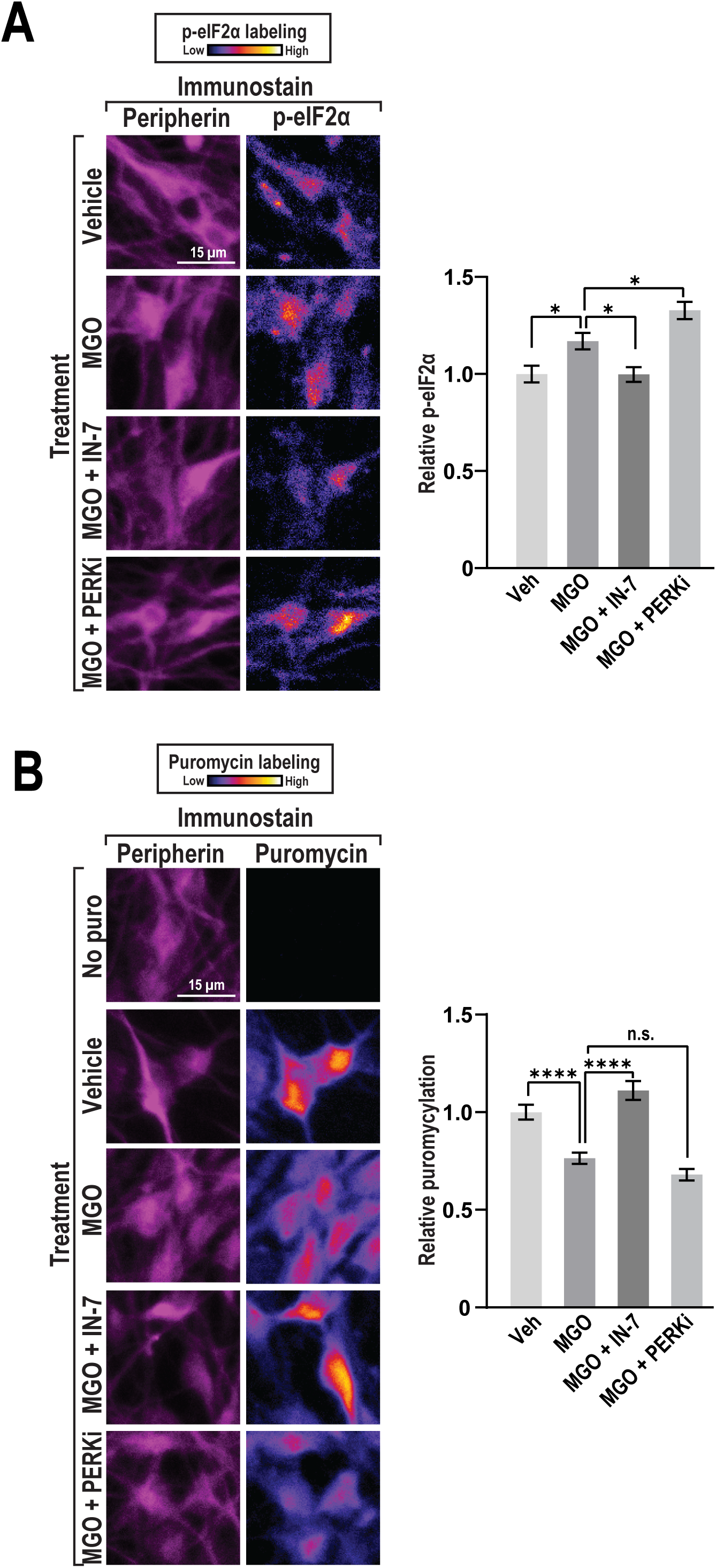
MGO induces the ISR through GCN2. Differentiated HD10.6 cells were treated for 24 hours with either vehicle, 1 µM MGO, 1 µM MGO with 100 nM GCN2-IN-7 (IN-7) added in the last hour of treatment, or 1 µM MGO with 50 nM GSK2606414 (PERKi) added in the last hour. (**A**) p-eIF2α CTCF was measured in peripherin-positive soma. Boxes represent mean p-eIF2α CTCF and bars depict ± SEM. (n = 114-125 cells per group), \**P* < 0.05; One-way ANOVA and Sidak’s multiple comparison test. Scale bar = 15 microns. (**B**) 1 µM puromycin was added in the last five minutes of treatment. Puromycin CTCF was measured in peripherin-positive soma. Boxes represent mean puromycin CTCF and bars depict ± SEM. (n = 108-115 cells per group), \*\*\*\**P* < 0.0001, n.s. = not significant; One-way ANOVA and Sidak’s multiple comparison test. Scale bar = 15 microns.

### MGO causes translational repression of ribosomal proteins in mice

To survey the effects of MGO on mRNA abundance and translation, ribosome profiling was conducted. The ribosome profiling method consists of isolation of ribosome-protected mRNA fragments followed by high-throughput sequencing [36] (**Fig. 2A**). As a control, total RNA is sequenced in parallel. Translation efficiency obtained by ribosome profiling is a more precise estimate of protein abundance than mRNA sequencing [47; 84]. This strategy has been applied to DRG tissues following nerve injury, a model of chemotherapy-induced neuropathy, and inflammatory mediators, but has yet to be applied to diabetic pain [7; 18; 20; 21; 49; 81]. Following systemic injection of MGO (2 mg/kg), we detected no significant changes in RNA abundance and over 1,000 changes in translation (**Fig. 2A**). Consistent with ISR activation, most transcripts with altered ribosome association exhibited reduced ribosome occupancy (1,050 total) while a subset of transcripts (51 total) had increased ribosome occupancy, suggestive of increased translational efficiency (**Fig. 2C**). Transcripts that exhibited increased ribosome occupancy were enriched for uORFs, consistent with their known roles in licensing mRNAs for translation during the ISR (**Supplementary Fig. 1**) [89]. Next, we asked which gene ontology terms were enriched among differentially translated mRNAs [15]. MGO triggered an increase in the translation of mRNAs linked to mechanical sensory perception, consistent with MGO causing mechanical pain through direct effects on sensory neurons [6; 55] (**Fig. 2D**). mRNAs linked to translation and ribosome biogenesis were repressed, suggestive of an inhibition of factors that mediate protein synthesis (**Fig. 2D**). To test this observation, we examined the effect of MGO on the abundance of newly translated ribosomal proteins in primary DRGs, as DRGs were used to conduct ribosome profiling (**Fig. 2E-F**). Newly translated proteins were labelled with puromycin, and samples were stained with nucleolin to visualize the nucleolus, as it is the site of ribosomal RNA transcription, processing, and subunit assembly. As a control, we inhibited the mechanistic target of rapamycin (mTOR) using Torin1, an ATP-competitive inhibitor of two distinct complexes that share the mTOR kinase (mTORC1 and mTORC2). We reasoned that blockade of mTOR would reduce translation in the nucleolus because it regulates nuclear import of ribosomal proteins and indirectly controls the availability of the cap-binding protein eIF4E [41; 74]. Addition of Torin1 diminished the abundance of newly translated proteins in the nucleolus as expected (**Fig. 2E-F**) (One-way ANOVA, F_2,314_ = 26.59, Sidak’s multiple comparison test, \*\*\*\**P* < 0.0001). Similarly, MGO reduced the presence of newly translated proteins in the nucleolus by 33%. This level of repression is similar to the repression of global translation seen previously (24%) (**Fig. 1B**). This observation is consistent with the notion that ribosomal mRNAs are among the most actively translated transcripts in the cell [78].

**Fig. 2.**
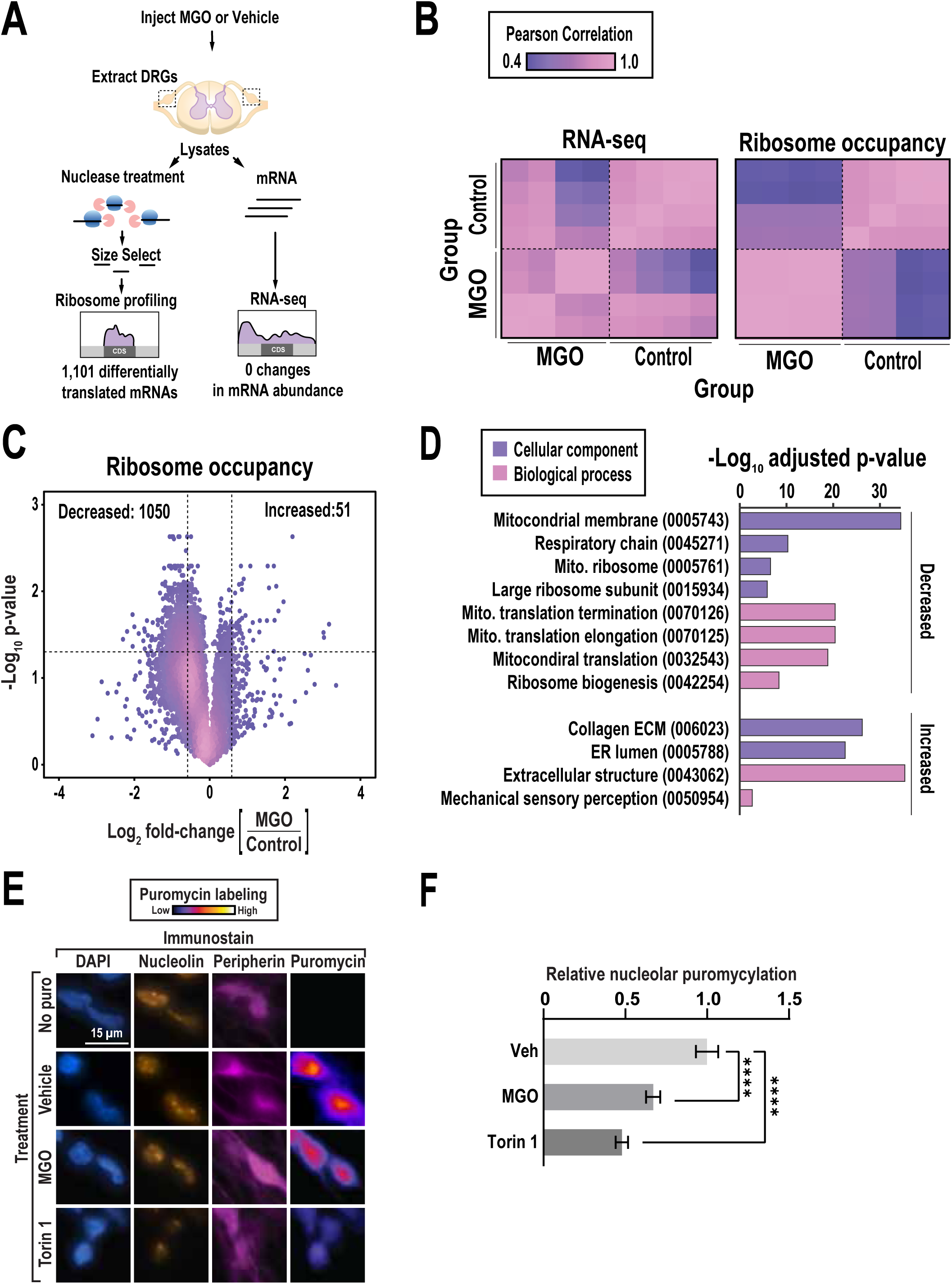
Ribosome profiling reveals broad impacts of MGO on protein synthesis. (**A**) Schematic depicting experimental setup. (**B**) Heatmap showing Pearson’s correlation comparing control and MGO groups from RNA-seq and ribosome profiling. (**C**) Volcano plot depicting the differential changes in translation. (**D**) Gene ontology analysis conducted with Enrichr. (**E**) Primary DRG neurons were cultured for six days and treated with either vehicle, 1 µM MGO for 24 hours, or 100 nM Torin1 for 30 minutes. 1 µM puromycin was added in the last ten minutes of treatment. Nucleolar puromycin CTCF was measured. (**F**) Quantification of nucleolar puromycin. Boxes represent mean puromycin CTCF and bars depict ± SEM. (n = 103-109 cells per group), \*\*\*\**P* < 0.0001; One-way ANOVA and Sidak’s multiple comparison test. Scale bar = 15 microns.

### Disruption of complex I of the electron transport chain triggers the ISR

One of the broad categories of mRNAs repressed by MGO comprises genes linked to the electron transport chain (**Fig. 2D**). This observation led us to ask if perturbation of the electron transport chain is upstream of GCN2. In cell lines, disruption of complex I activates the ISR in a GCN2-dependent manner through a mechanism involving amino acid depletion [57]. To determine if complex I inhibition results in GCN2 activation in neurons, HD10.6 cells were differentiated and treated with the small-molecule inhibitor piericidin A, which increased p-eIF2α by 32% (**Fig. 3A**) (One-way ANOVA, F_3,445_ = 58.07, Sidak’s multiple comparison test, \*\**P* < 0.01, \*\*\**P* < 0.001, \*\*\*\**P* < 0.0001). Co-treatment with a GCN2 inhibitor fully reversed this effect (**Fig. 3A**). We conclude that GCN2 is activated in response to complex I inhibition in sensory neurons.

**Fig. 3.**
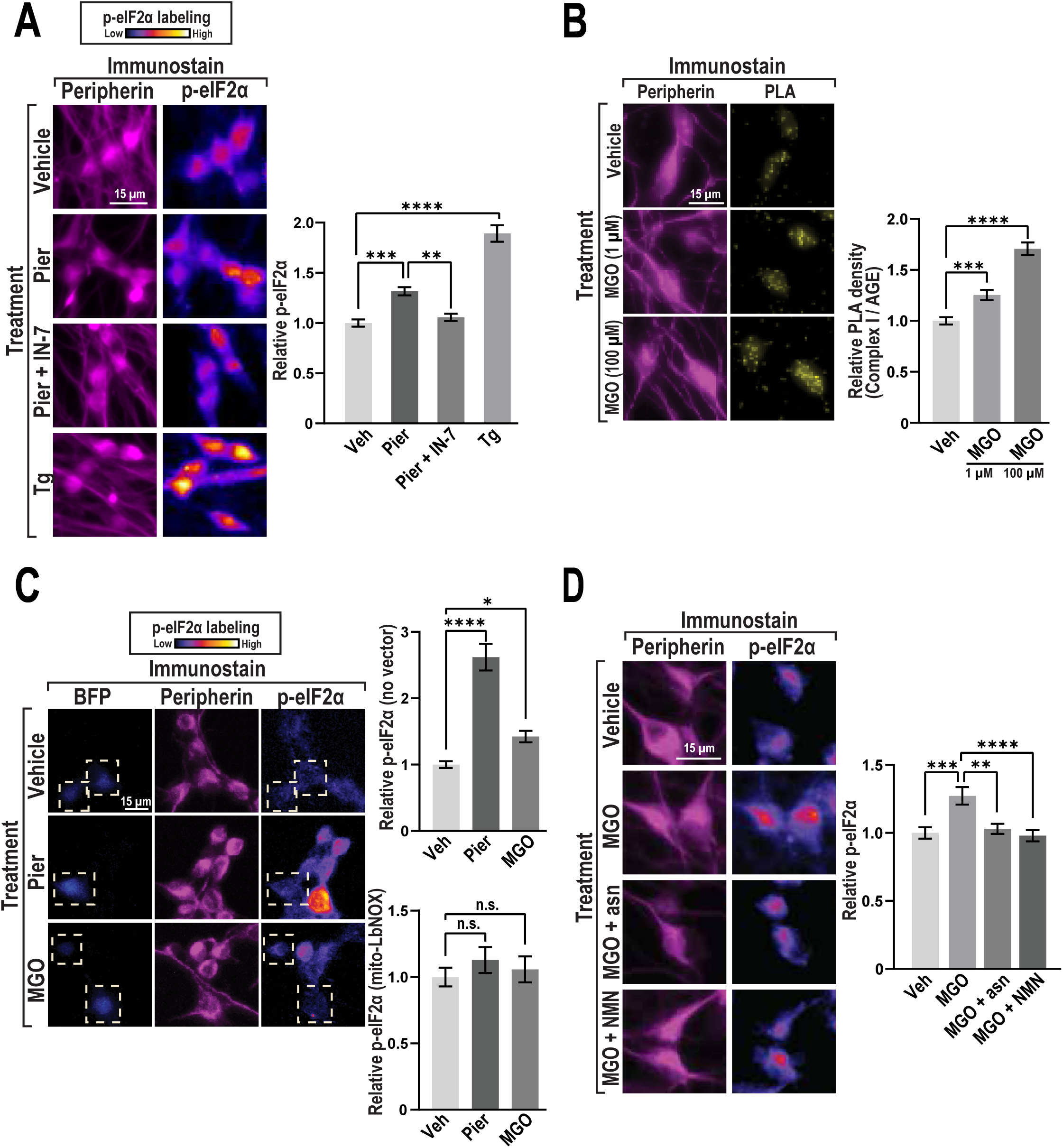
Complex I inhibition induces the ISR through GCN2. **(A)** Differentiated HD10.6 cells were treated with either vehicle, 500 nM piericidin A for six hours, 500 nM piericidin A with 100 nM IN-7 for six hours, or 1 µM thapsigargin (Tg) for 30 minutes. p-eIF2α CTCF was measured in peripherin-positive cells. Boxes represent mean p-eIF2α CTCF and bars depict ± SEM. (n = 108-116 cells per group), \*\**P* < 0.01, \*\*\**P* < 0.001, \*\*\*\**P* < 0.0001; One-way ANOVA and Sidak’s multiple comparison test. Scale bar = 15 microns. (**B**) Differentiated HD10.6 cells were treated for 24 hours with vehicle, 1 µM MGO, or 100 µM MGO. PLA assay was performed with antibodies targeting advanced glycation end products (AGE) and complex I. PLA signal was measured in peripherin-positive cells. Boxes represent mean PLA puncta density and bars depict ± SEM. (n = 119-123 cells per group), \*\*\**P* < 0.001, \*\*\*\**P* < 0.0001; One-way ANOVA and Sidak’s multiple comparison test. Scale bar = 15 microns. (**C**) F11 cells were transfected with either control or mitoLbNOX and differentiated prior to treatments. Cells successfully transfected with mitoLbNOX were identified by measuring blue fluorescent protein (BFP) signal. Cells were treated with either vehicle, 1 µM MGO for 24 hours, or 500 nM piericidin A for six hours. p-eIF2α CTCF was measured in peripherin-positive BFP^-^ cells and peripherin-positive BFP^+^ cells. Boxes represent mean p-eIF2α CTCF and bars depict ± SEM. (n = 71-118), n.s. = not significant, \**P* < 0.05, \*\*\*\**P* < 0.0001; One-way ANOVA and Sidak’s multiple comparison test. Scale bar = 15 microns. (**D**) Differentiated HD10.6 cells were treated with either vehicle, 1 µM MGO, 1 µM MGO with 500 µM asparagine, or 1 µM MGO with 100 µM nicotinamide mononucleotide (NMN) for 24 hours. p-eIF2α CTCF was measured in peripherin-positive cells. Boxes represent mean p-eIF2α CTCF and bars depict ± SEM. (n = 90 cells per group), \*\**P* < 0.01, \*\*\**P* < 0.001, \*\*\*\**P* < 0.0001; One-way ANOVA and Sidak’s multiple comparison test. Scale bar = 15 microns.

A critical question is whether MGO impacts NAD redox state. As a positive control, we confirmed that inhibition of complex I with piericidin A increased the ratio of NADH/NAD^+^ (**Supplementary Fig. 2A**). However, the sensitivity of these measurements is limited as lower concentrations of piericidin A failed to evoke detectable changes (**Supplementary Fig. 2A**) (One-way ANOVA, F_4,10_ = 4.981, Sidak’s multiple comparison test, n.s. = not significant, \**P* < 0.05, \*\**P* < 0.01). Similarly, high doses of MGO also resulted in increased levels of NADH/NAD^+^ while low doses did not (**Supplementary Fig. 2B**) (One-way ANOVA, F_5,12_ = 78.25, Sidak’s multiple comparison test, n.s. = not significant, \*\*\*\**P* < 0.0001). We next examined how MGO impacts complex I. Given that MGO reacts directly with proteins to form advanced glycation end products, we asked if MGO forms advanced glycation end products on complex I using proximity ligation assays [46]. This approach enables hybridization of fluorescent probes to a template formed only when two antigens are within 40 nm of each other [73]. Using antibodies against complex I and advanced glycation end products, we found that MGO resulted in an increase in PLA signal at both a low and a high dose in differentiated HD10.6 cells (**Fig. 3B**) (One-way ANOVA, F_2,359_ = 50.30, Sidak’s multiple comparison test, \*\*\**P* < 0.001, \*\*\*\**P* < 0.0001). Impaired complex I can impact downstream electron transport chain complexes, including ATP production through ATP synthase. Piericidin A lowers the ATP/ADP ratio in cell lines [57]. We tested whether Piericidin A and MGO lead to changes in cellular ATP levels. We blocked glycolysis with a small molecule inhibitor to prevent compensation from other metabolic pathways that would mask differences. We found that both piericidin A and MGO at two concentrations lower ATP levels in human sensory neurons (**Supplementary Fig. 2C**) (One-way ANOVA, F_3,8 =_ 7.570, Dunnett’s multiple comparison test, \**P* < 0.05, \*\**P* < 0.01). These data indicate that MGO reacts with complex I, increases the ratio of NADH/NAD^+^, and lowers ATP levels. This raises a logical question – does the imbalance in NADH/NAD^+^ cause GCN2 activation by MGO? To test this idea, we made use of a genetic tool consisting of a mitochondrial localization sequence and a bacterial NADH oxidase called LbNOX [79]. We used differentiated F11 cells as they are more amenable to transfection compared to HD10.6 cells. Transfection of this construct, termed mitoLbNOX, prevented induction of p-eIF2α by either MGO or piericidin A (**Fig. 3C**) (One-way ANOVA, F_2,269_ = 0.8209, Sidak’s multiple comparison test, n.s. = not significant) (One-way ANOVA, F_2,297_ = 45.27, Sidak’s multiple comparison test, \**P* < 0.05, \*\*\*\**P* < 0.0001). This suggests that oxidation of the mitochondrial pool of NADH prevents ISR activation due to either complex I inhibition or addition of MGO (**Fig. 3C**). Downstream of NAD^+^ metabolism is asparagine synthesis [57]. We tested whether supplementing with asparagine or a precursor to NAD^+^, nicotinamide mononucleotide (NMN), prevents activation of the ISR by MGO. We found that both asparagine and NMN prevented ISR induction by MGO in differentiated HD10.6 cells (**Fig. 3D**) (One-way ANOVA, F_3,355_ = 8.261, Sidak’s multiple comparison test, \*\**P* < 0.01, \*\*\**P* < 0.001, \*\*\*\**P* < 0.0001). This suggests that asparagine depletion downstream of complex I inhibition drives ISR activation.

### GCN2 deletion prevents neuronal sensitization by a TRPA1 agonist

To understand the role of GCN2 in a diabetic animal, we utilized the streptozotocin model of type I diabetes (T1D). Mice were administered streptozotocin to ablate pancreatic beta cells, and blood glucose levels were monitored to confirm diabetes [31]. We measured neuronal sensitization, as it is a defining feature of neuropathic pain. Both MGO and streptozotocin lead to sensitization of sensory neurons [3; 4; 6; 34; 55]. Our prior work showed that inhibiting GCN2 with IN-7 prevented MGO-induced sensitization [55]. However, it is unknown whether GCN2 plays a role in neuronal sensitization in the context of T1D evoked by streptozotocin treatment. To test this, we performed calcium imaging in either vehicle or streptozotocin injected *Pirt*^Cre/+^ mice. To determine the role of GCN2, streptozotocin was injected into *GCN2*^fl/fl^ *Pirt*^Cre/+^ mice. We stimulated neurons with a TRPA1 agonist, as streptozotocin indirectly promotes neuronal sensitization in a TRPA1-dependent manner [3; 34]. We found that streptozotocin resulted in a significant increase in the magnitude of calcium influx evoked by activating TRPA1 (**Fig. 4**). Genetic deletion of GCN2 prevented this effect (**Fig. 4**). (One-way ANOVA, F_2,239_ = 4.243, Sidak’s multiple comparison test, \**P* < 0.05). This suggests that streptozotocin sensitizes TRPA1 channels via GCN2.

**Figure 4.**
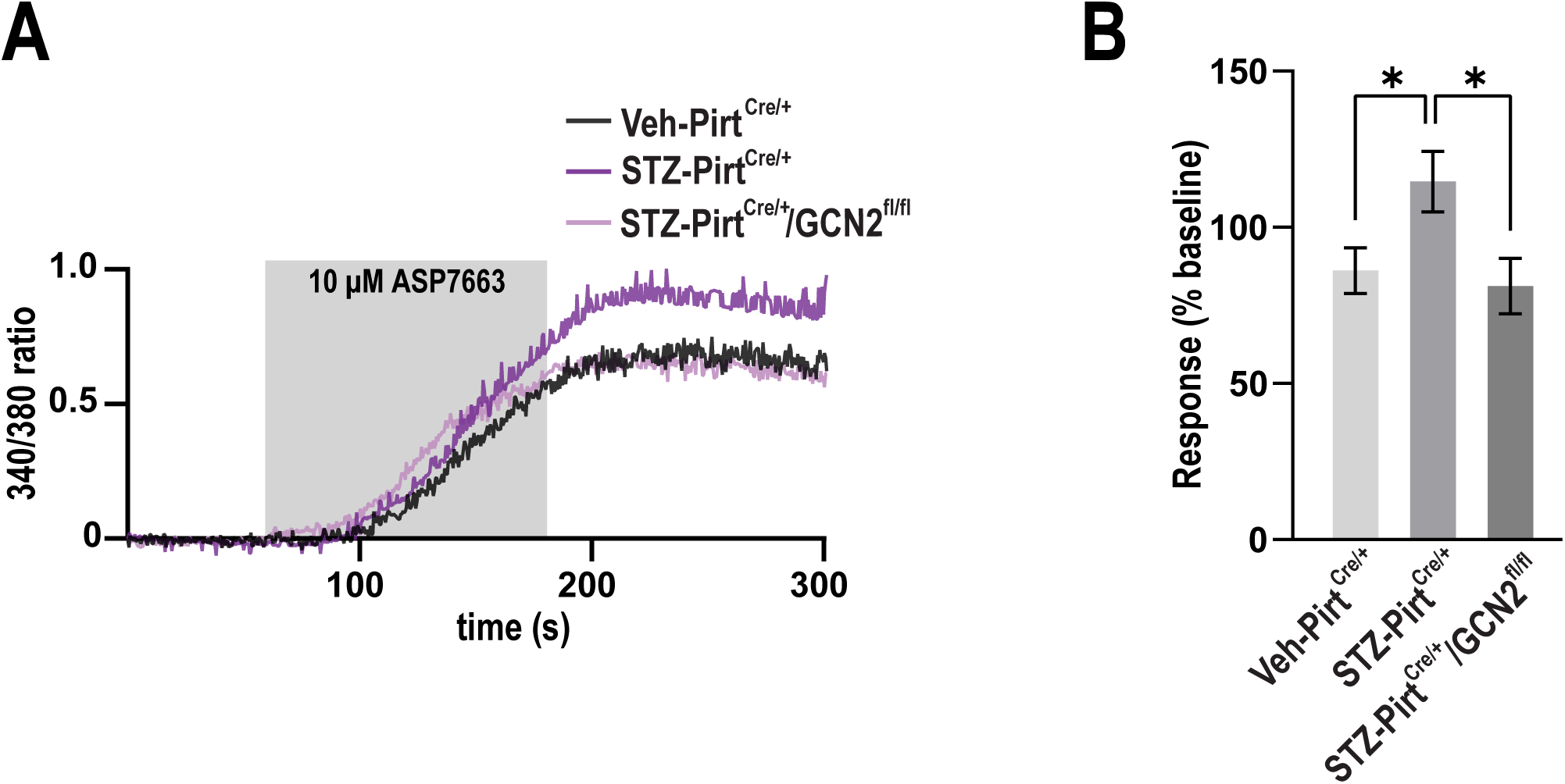
GCN2 deletion prevents STZ-induced TRPA1 sensitization. (A) Composite calcium imaging trace from DRG neurons isolated from either vehicle or STZ-treated *Pirt*^Cre/+^ mice or STZ-treated *GCN2*^fl/fl^ *Pirt*^Cre/+^ mice four weeks following administration of STZ. 10 µM ASP7663 was applied to cells and their activity was measured. (B) The quantification of the traces represents the response magnitude (% increase over baseline after ASP7663 stimulus. Boxes represent the mean response magnitude, bars represent ± SEM. (n = 75-84 cells per group). \**P* < 0.05; One-way ANOVA and Sidak’s multiple comparison test.

### GCN2 is activated in a model of type I diabetes

We next examined if GCN2 is required for ISR activation *in vivo* in the streptozotocin model. To measure mechanical allodynia, we assayed withdrawal thresholds using calibrated von Frey filaments. Withdrawal thresholds were significantly lower one month after streptozotocin treatment in both male and female mice relative to vehicle injection, indicative of sensitization (**Fig. 5A and 5C**) (Two-way ANOVA, F_1,13_ = 114.0, Bonferroni’s multiple comparison test, \*\*\*\**P* < 0.0001) (Two-way ANOVA, F_1,14_ = 223.5, Bonferroni’s multiple comparison test, \**P* < 0.05, \*\*\**P* < 0.001, \*\*\*\**P* < 0.0001). Mice were then treated with a GCN2 inhibitor or a vehicle control. Animals that received the inhibitor returned to baseline transiently in both males and females (**Fig. 5B and 5D**) (Two-way ANOVA, F_1,84_ = 136.2, Bonferroni’s multiple comparison test, \*\**P* < 0.01, \*\*\*\**P* < 0.0001) (Two-way ANOVA, F_1,84_ = 99.13, Bonferroni’s multiple comparison test, \*\*\**P* < 0.001, \*\*\*\**P* < 0.0001). As a more precise genetic test, GCN2 was ablated specifically in sensory neurons by crossing pan-sensory neuron *Pirt*^Cre/Cre^ with a GCN2 exon XII-flox strain. GCN2 levels were significantly reduced in DRG tissues obtained from mutant animals (**Supplementary Fig. 3**) (Unpaired t-test with Welch’s correction, P value = 0.0165, two-tailed, \**P* < 0.05). Conditional elimination of GCN2 significantly diminished mechanical allodynia in animals treated with streptozotocin (**Fig. 5E and 5F**) (Two-way ANOVA, F_1,13_ = 25.23, Bonferroni’s multiple comparison test, \**P* < 0.05, \*\**P* < 0.01, \*\*\*\**P* < 0.0001) and (Two-way ANOVA, F_1,12_ = 57.68, Bonferroni’s multiple comparison test, \**P* < 0.05, \*\*\**P* < 0.001). To determine if GCN2 is required for ISR induction in DRG tissues, we examined p-eIF2α levels in DRG tissues (**Fig. 6A**). We compared two classes of sensory neurons using established markers, peptidergic (TRPV1) and non-peptidergic (IB4). Streptozotocin resulted in strong ISR induction in TRPV1^+^ neurons but not IB4^+^ neurons (**Fig. 6B and 6C**) (One-way ANOVA, F_2,9_ = 32.34, Bonferroni’s multiple comparison test, \*\**P* < 0.01, \*\*\*\**P* < 0.0001) (One-way ANOVA, F_2,9_ = 0.9931, Bonferroni’s multiple comparison test, n.s. = not significant). Elimination of GCN2 reduced ISR induction but did not fully reduce p-eIF2α to basal levels, suggestive of either functional redundancy among ISR kinases, inhibition of the eIF2α phosphatase (CReP), or incomplete penetrance from the Cre-lox system. We conclude that GCN2 function in sensory neurons is critical for ISR induction and mechanical sensitivity in a diabetic mouse model.

**Fig. 5.**
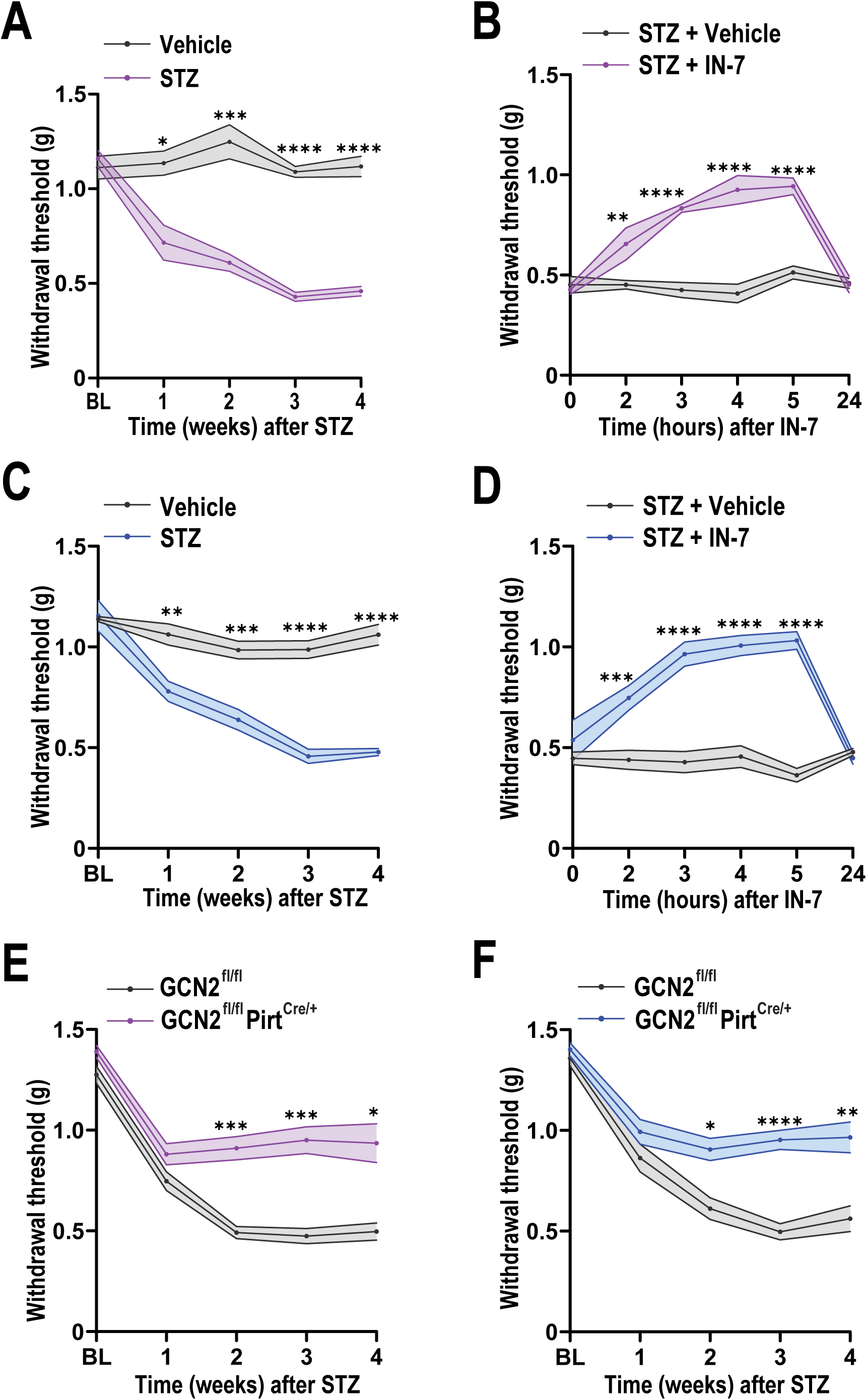
GCN2 inhibition alleviates STZ-induced mechanical sensitization. (**A**) Female WT B6 mice were injected with either vehicle or STZ (50 mg/kg, i.p.) for five consecutive days. Von Frey was performed to determine withdrawal threshold every week for four weeks following STZ injections. Lines represent mean and the shaded area depicts ± SEM (n = 8 animals per group). \**P* < 0.05, \*\*\**P* < 0.001, \*\*\*\**P* < 0.0001; Two-way ANOVA with Bonferroni’s multiple comparisons test. (**B**) Female STZ mice were injected with either vehicle or IN-7 (5 mg/kg) and von Frey was performed one-hour post-injection through 24 hours. Lines represent mean and the shaded area depicts ± SEM (n = 8 animals per group). \*\**P* < 0.01, \*\*\*\**P* < 0.0001; Two-way ANOVA with Bonferroni’s multiple comparisons test. (**C**) Male WT B6 mice were injected with either vehicle or STZ (50 mg/kg, i.p.) for five consecutive days. Von Frey was performed to determine withdrawal threshold every week for four weeks following STZ injections. Lines represent mean and the shaded area depicts ± SEM (n = 7-8 animals per group). \*\**P* < 0.01, \*\*\**P* < 0.001, \*\*\*\**P* < 0.0001; Two-way ANOVA with Bonferroni’s multiple comparisons test. (**D**) Male STZ mice were injected with either vehicle or IN-7 (5 mg/kg) and von Frey was performed one-hour post-injection through 24 hours. Lines represent mean and the shaded area depicts ± SEM (n = 8). \*\*\**P* < 0.001, \*\*\*\**P* < 0.0001; Two-way ANOVA with Bonferroni’s multiple comparisons test. (**E**) STZ (50 mg/kg, i.p.) was administered to either female *GCN2*^fl/fl^ or female *GCN2*^fl/fl^ *Pirt*^Cre/+^ animals. Von Frey was performed to determine withdrawal threshold every week for four weeks following STZ injections. Lines represent mean and the shaded area depicts ± SEM (n = 7 animals per group). \**P* < 0.05, \*\*\**P* < 0.001; Two-way ANOVA with Bonferroni’s multiple comparisons test. (**F**) STZ (50 mg/kg, i.p.) was administered to either male *GCN2*^fl/fl^ or male *GCN2*^fl/fl^ *Pirt*^Cre/+^ animals. Von Frey was performed to determine withdrawal threshold every week for four weeks following STZ injections. Lines represent mean and the shaded area depicts ± SEM (n = 7-8 animals per group). \**P* < 0.05, \*\**P* < 0.01, \*\*\*\**P* < 0.0001; Two-way ANOVA with Bonferroni’s multiple comparisons test.

**Fig. 6.**
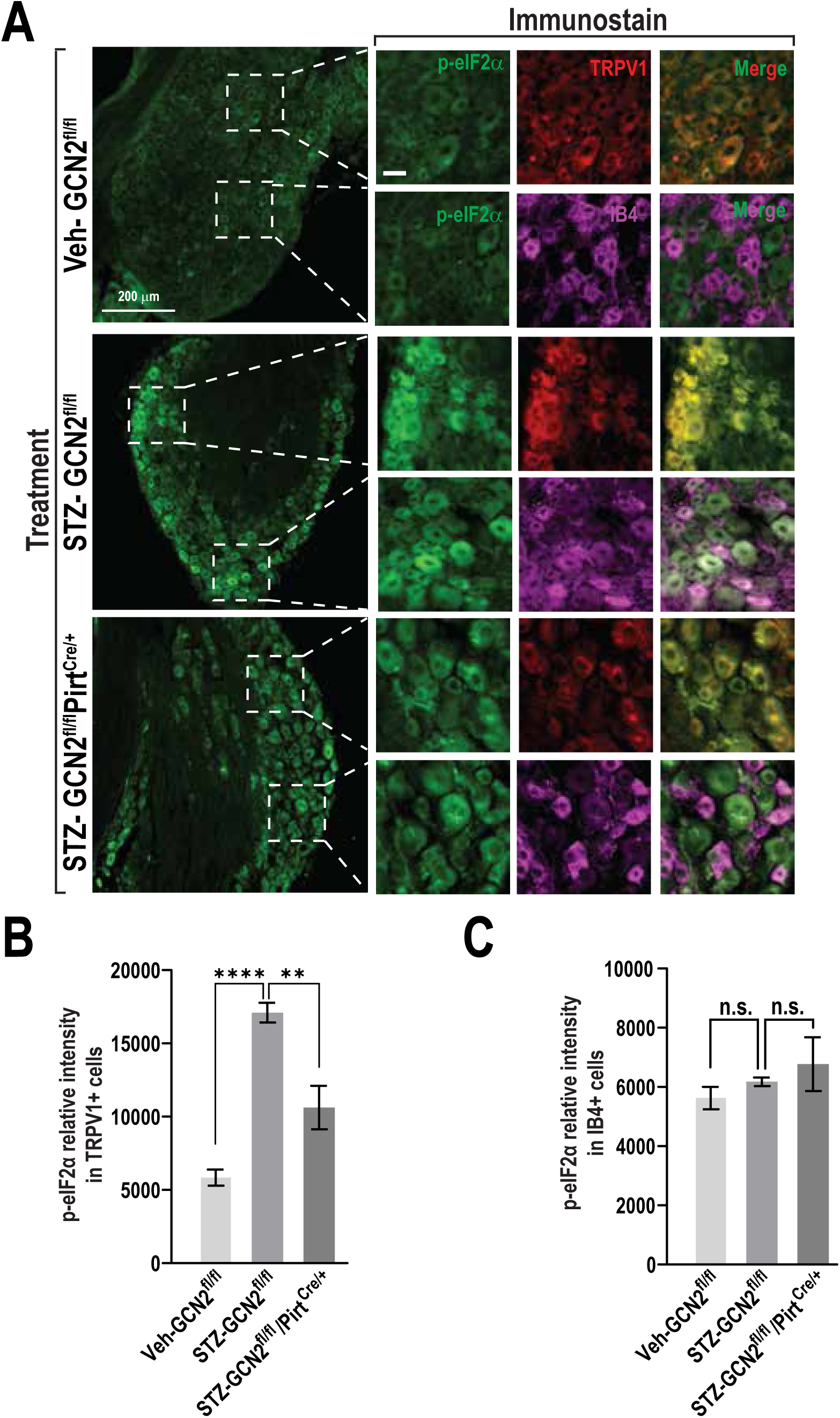
STZ induces the ISR in TRPV1^+^ neurons through GCN2 *in vivo*. (**A**) Immunohistochemistry staining showing p-eIF2α expression in transient receptor potential vanilloid 1 (TRPV1) and isolectin B4 (IB4) neurons/cells. *GCN2*^fl/fl^ animals were injected with either vehicle or STZ (50 mg/kg, i.p.). *GCN2*^fl/fl^ *Pirt*^Cre/+^ mice were injected with STZ (50 mg/kg, i.p.). Four weeks following vehicle or STZ injections, DRG tissues were harvested and stained for TRPV1, IB4, and p-eIF2α. (**B**) Quantification of p-eIF2α fluorescence intensity in TRPV1^+^ cells. Boxes represent mean fluorescence intensity and bars depict ± SEM. (n = 4 mice per group), \*\**P* < 0.01, \*\*\*\**P* < 0.0001. C) Quantification of p-eIF2α intensity in IB4^+^ cells. Boxes represent mean fluorescence intensity and bars depict ± SEM. (n = 4 mice per group). n.s. = not significant. One-way ANOVA with Bonferroni’s multiple comparisons test.

### Defective electron transport leads to GCN2-dependent mechanical allodynia that can be reversed by metabolic reprogramming

In human sensory neurons, we observed that piericidin A induces the ISR in a GCN2-dependent manner (**Fig. 3A**). We then wanted to determine whether complex I inhibition is sufficient to cause mechanical allodynia and if GCN2 deletion would prevent sensitization. To test this, piericidin A was injected into either *GCN2*^fl/fl^ or *GCN2*^fl/fl^ *Pirt*^Cre/+^ mice. Piericidin A leads to a significant decrease in the withdrawal threshold three hours after injection in *GCN2*^fl/fl^ mice (**Fig. 7A**) (Two-way ANOVA, F_1,17_ = 32.31, Bonferroni’s multiple comparison test, \*\**P* < 0.01, \*\*\**P* < 0.001, \*\*\*\**P* < 0.0001). However, conditional deletion of GCN2 significantly diminished mechanical allodynia and these mice returned to baseline sooner than *GCN2*^fl/fl^ mice. (**Fig. 7A**) This suggests that complex I inhibition is sufficient to drive mechanical sensitization and that this is prevented by GCN2 deletion.

**Fig. 7.**
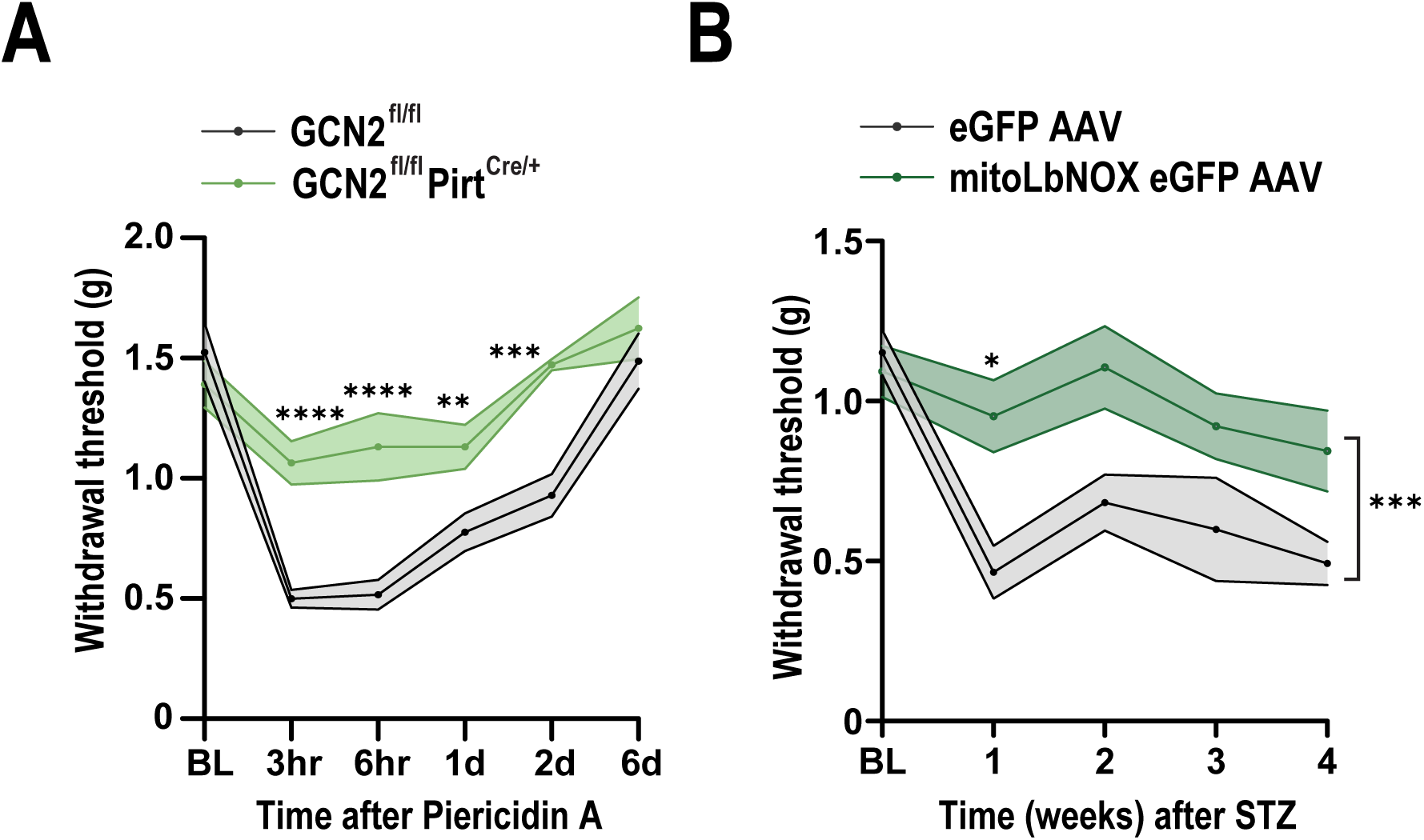
Defective electron transport leads to GCN2-dependent mechanical allodynia that can be reversed by mitoLbNOX. (**A**) *GCN2*^fl/fl^ or *GCN2*^fl/fl^ *Pirt*^Cre/+^ mice were injected with piericidin A (0.1 µg, i.pl.). Von Frey was performed to determine withdrawal threshold up to six days post injection. Lines represent mean and the shaded area depicts ± SEM (n = 9-10 animals per group). \*\**P* < 0.01, \*\*\**P* < 0.001, \*\*\*\**P* < 0.0001; Two-way ANOVA with Bonferroni’s multiple comparisons test. (**B**) *Pirt*^Cre/+^ mice were injected intrathecally with either the control or mitoLbNOX AAV. Two weeks following AAV injection, STZ (50 mg/kg, i.p.) was injected into both groups. Von Frey was performed to determine withdrawal threshold every week for four weeks following STZ injections. Lines represent mean and the shaded area depicts ± SEM (n = 10 animals per group). \**P* < 0.05, \*\*\**P* < 0.001; Two-way ANOVA with Bonferroni’s multiple comparisons test.

To determine if oxidation of NADH in the mitochondria prevents mechanical allodynia *in vivo*, we injected mice intrathecally with either mitoLbNOX or a GFP control packaged in AAV-MaCPNS1, which has tropism for DRG sensory neurons [12]. Two weeks after administration of the AAVs, we administered streptozotocin and measured mechanical withdrawal thresholds. Mice injected with mitoLbNOX exhibited significantly reduced sensitivity one week after treatment with streptozotocin (**Fig. 7B**) (Two-way ANOVA, F_1,18_ = 22.76, Bonferroni’s multiple comparison test, \**P* < 0.05). This group was less sensitive than control mice for the following three weeks. The data indicate that metabolic reprogramming partially reduces sensitization in a diabetic mouse model. Collectively, these observations reveal that MGO disrupts complex I through the formation of advanced glycation end products (**Fig. 8**). This results in depletion of NAD^+^ and asparagine. Metabolic dysfunction consequently drives ribosome collisions, GCN2 activation, and mechanical hypersensitivity.

**Fig. 8.**
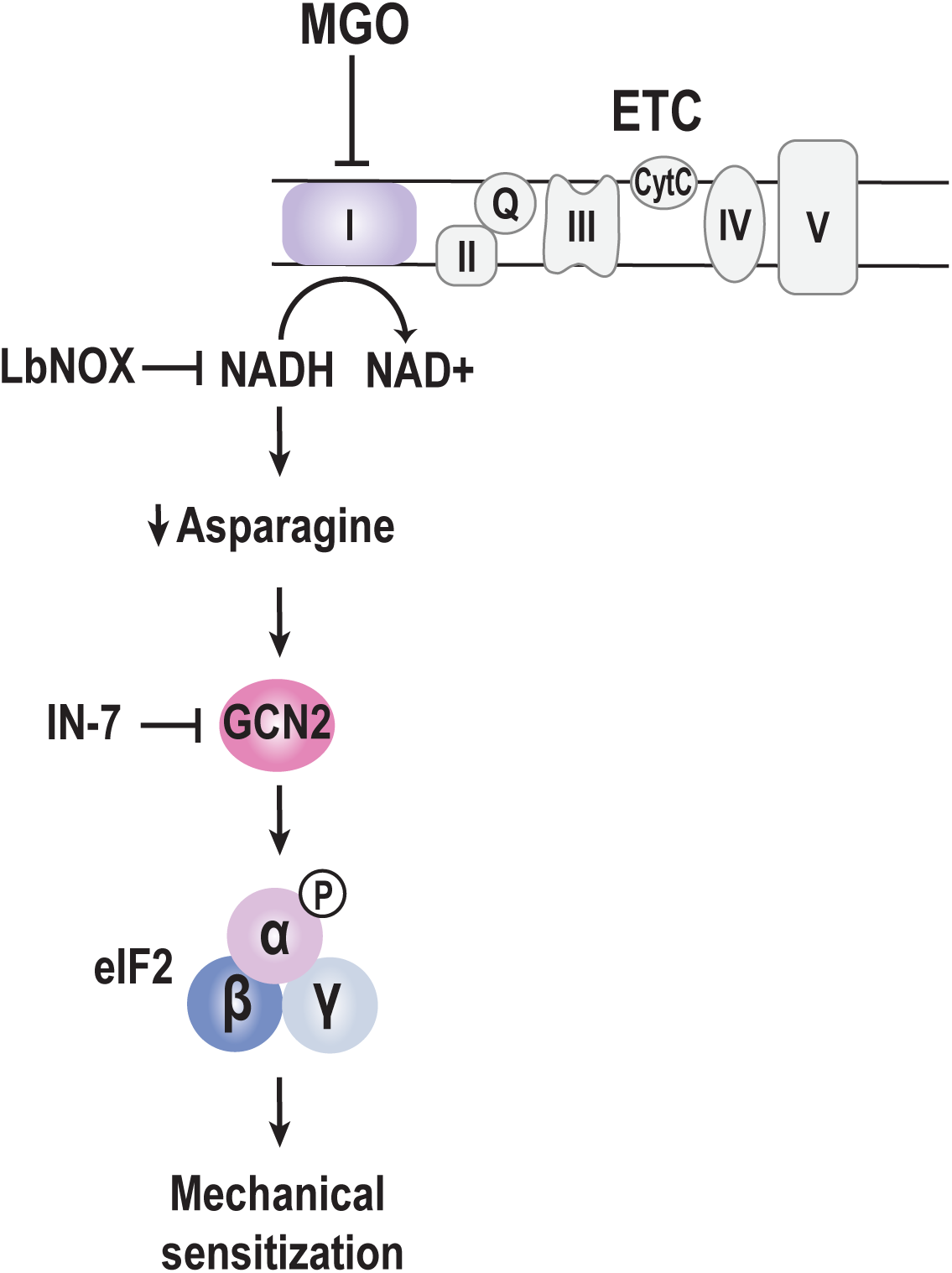
A proposed model for MGO’s mechanism of action. MGO causes complex I dysfunction, leading to an increased NADH/NAD^+^ ratio. Decreased NAD^+^ leads to decreased asparagine synthesis. This promotes GCN2-dependent ISR induction via eIF2α phosphorylation and causes mechanical sensitization.

## Discussion

Our findings enable five major conclusions. First, we establish GCN2 as the conserved mediator of ISR activation and translational repression downstream of MGO in human sensory neurons. Second, we demonstrate that MGO causes widespread translational repression *in vivo*, selectively targeting highly translated transcripts encoding ribosomal components and electron transport chain machinery. Third, we show that MGO directly glycates complex I and disrupts mitochondrial NAD redox homeostasis (NADH/NAD⁺ ratio) to trigger GCN2 activation. Fourth, metabolic reprogramming through the introduction of a bacterial mitochondrial NADH oxidase prevents ISR induction by MGO and significantly reduces streptozotocin-induced mechanical allodynia. Fifth and finally, genetic or pharmacological disruption of GCN2 blocks activation of the ISR and alleviates mechanical allodynia in a model of diabetic neuropathy. Below, we discuss each finding and its broader implications.

The ISR acts as a central node in the regulation of neuronal excitability and pain, but the upstream kinases that detect metabolic insults are unclear in human sensory neurons [6; 19; 42; 55; 58; 70]. While previous reports have implicated both PERK and GCN2 as downstream of MGO in rodents, our data clearly indicate that MGO-induced eIF2α phosphorylation and global translational suppression are dependent on GCN2 in human sensory neurons [6; 55]. This solidifies GCN2 as the principal sensor of MGO-mediated stress across species. Importantly, activation of GCN2 is sufficient to cause mechanical allodynia in rodents [58]. Intraplantar injection of halofuginone, a GCN2 activator, leads to a significant decrease in withdrawal thresholds. However, mice lacking GCN2 are significantly less sensitive to halofuginone [58]. These findings emphasize that GCN2 and ISR activation are sufficient to drive pain *in vivo*. Notably, GCN2 is increasingly recognized as a general sentinel of translational and metabolic stress in an array of disease states [59]. Impairments in elongation, whether triggered by defective tRNA charging in motor neurons (Charcot-Marie-Tooth disease and paclitaxel-induced neuropathy), ribosome collisions in striatal neurons (Huntington’s disease), or eEF2K-mediated phosphorylation downstream of inflammation in sensory neurons, all converge on GCN2 activation as a major driver of pathological signaling [5; 54; 58; 70; 75]. Our findings extend this paradigm to painful diabetic neuropathy.

By leveraging unbiased ribosome profiling *in vivo*, we discovered a striking divergence in how MGO differentially impacts transcriptional and translational controls. While transcription – as judged by RNA abundance – remained largely unaltered, MGO induced profound alterations in protein synthesis. This was evidenced by repression of transcripts encoding ribosomal proteins and structural components of the electron transport chain. Ribosomal mRNAs are among the most actively translated messages in the cell. Their repression by MGO likely reflects an energy-conservation response elicited by translational stress. Paradoxically, MGO selectively stimulated the translation of a subset of transcripts linked to mechanical sensory perception. This dual action is a hallmark of the ISR, where global translation attenuation coexists with the preferential translation of stress-adaptive mRNAs containing uORFs [33; 52; 82]. These regulatory elements were highly enriched in mRNAs with increased translation in DRG. An important next question asks which mRNAs are responsible for driving the maladaptive plasticity seen in response to engagement of the ISR. Our prior work identified brain-derived neurotrophic factor (BDNF) as a functional downstream target of GCN2 [70]. However, BDNF is largely dispensable for neuropathic pain [22]. Thus, a critical problem moving forward is the identification of the relevant pronociceptive mRNA(s) whose translation promotes pain amplification.

How does MGO reprogram protein synthesis? Our study uncovers a translational signaling axis downstream of mitochondrial complex I. Using a proximity ligation assay approach, we demonstrated that MGO directly glycates complex I, forming advanced glycation end products. Glyoxalases are enzymes that detoxify MGO to prevent cellular damage due to glycation events [32]. Importantly, reduced glyoxalase activity has been implicated in diabetic pain in patients [4; 9; 38; 69]. Interestingly, advanced glycation end products are also implicated in diabetic complications [11; 83]. This mechanism could be shared in other diseases, as MGO impairs complex I in both malignant cancer cells and cardiac cells [29]. Complex I function is essential for regenerating the NAD⁺ pool required for upstream amino acid catabolism and metabolic recycling; its disruption leads to localized amino acid starvation, uncharged tRNAs, and subsequent GCN2 activation [57]. Reduced NAD^+^ has been observed in diabetic tissues [25]. NAD^+^ precursor supplementation has been shown to reverse diabetic peripheral neuropathy in animal models [13; 80]. Crucially, correcting this redox imbalance using the genetically encoded mitochondrial NADH oxidase mitoLbNOX blocked ISR activation induced by either MGO or the complex I inhibitor piericidin A. Similarly, addition of NMN also prevented ISR induction, which further supports the notion that NAD^+^ metabolism is downstream of MGO. This demonstrates that mitochondrial redox stress is not merely a byproduct of diabetic metabolism, but an active, causative trigger of GCN2-dependent translational control in nociceptors. A not mutually exclusive hypothesis is that MGO activates the ISR through energy depletion. Because translation requires high amounts of energy and MGO reduces ATP levels, this energy deficit may contribute to ISR activation. In cell lines, ATP depletion through inhibition of ATP synthase also leads to ISR activation [57].

A central question highlighted by our work is whether genetic reprogramming of metabolism in sensory neurons can be leveraged to diminish pain-associated behaviors. By normalizing the NADH/NAD⁺ balance with mitoLbNOX, ISR engagement was prevented and streptozotocin-induced mechanical allodynia was significantly diminished *in vivo*. This suggests that targeting metabolic-translational control crosstalk in peripheral sensory neurons can prevent established neuropathic behaviors without requiring central nervous system penetration. What is the translational significance of this finding? Given that impairment of complex I drives asparagine depletion, we also demonstrated that asparagine supplementation prevents ISR activation by MGO. Reduced levels of asparagine occur as a consequence of diabetes [40]. The observation that asparagine supplementation prevents activation of the ISR by MGO implies that asparagine depletion is the proximal driver of ribosome collisions.

We found that mitoLbNOX does not fully diminish mechanical sensitization *in vivo*. There are multiple potential explanations as to why the rescue is incomplete. One relates to the diverse effects of MGO. In addition to proteins, MGO also modifies RNA, and these RNA adducts are elevated in type 2 diabetes [51]. RNA modifications are known to impede elongation, and our prior work demonstrated that MGO triggers ribosome collisions in sensory neurons [55; 68]. Thus, while MGO impairs complex I, it also likely glycates mRNA, leading to ribosome slowdown and collisions. However, the finding that ISR activation is largely reversed upon expression of mitoLbNOX suggests that metabolism is the major driver of GCN2 activation.

A limitation of our animal model is that it does not fully reproduce the effects of diabetes on sensory neurons in patients. For example, the streptozotocin mouse model does precisely replicate the progressive neuronal loss observed in human diabetic neuropathy that occurs over years and/or decades [26; 37]. An additional limitation is that we focused on evoked measurements of mechanical hypersensitivity as the behavioral endpoint throughout this work. Moving forward, an important question is if affective dimensions of pain also require GCN2 in sensory neurons.

In summary, this work identifies GCN2 as a conserved bridge linking MGO-induced glycolytic stress, mitochondrial complex I glycation, and translational dysregulation in diabetic neuropathic pain. Mitochondrial NADH oxidation and disruption of GCN2 rescue translational homeostasis and alleviate mechanical hypersensitivity. Targeting the GCN2 signaling axis represents a promising strategy for reversing pain amplification as a consequence of metabolic dysfunction.

## Ethics statement

All animal procedures were approved by the Institutional Animal Care and Use Committee (protocol ID: M006582-R01-A06) at the University of Wisconsin-Madison.

## AI disclosure statement

Gemini was used to edit the manuscript. The authors take full responsibility for the content and accuracy of the final document.

## Acknowledgments

We thank Dr. June Bryan de la Peña and Tzu-Fang (Cherry) Lou for assistance with sample preparation.

## Author contributions

Conceptualization: A.R.M and Z.T.C.

Methodology: A.R.M and Z.T.C.

Investigation: A.R.M (both of them), G.G., S.R., J.B.S, and R.C

Supervision: Z.T.C.

Writing—original draft: A.R.M and Z.T.C.

Writing—review & editing: A.R.M (both of them), G.G., S.R., J.B.S, and R.C

## Data, code, and materials availability

Sequencing data are available through GEO tokens: GSE230457 and GSE230458.

## Figure Legends

**Fig. S1. MGO upregulates the translation of uORF-containing transcripts** Cumulative fraction plot showing the number of uORFs (MGO RFP # uORFs) across differentially expressed gene subsets compared to all evaluated genes. Genes upregulated upon MGO treatment (Up, green line; n = 71) show a significantly higher proportion of transcripts with multiple uORFs compared to background (All genes, black line). Genes downregulated upon treatment (Down, red line; n = 2,123) contain fewer uORFs relative to background. P values were obtained from Kolmogorov-Smirnov tests.

**Fig. S2.** Piericidin A and MGO alter NAD/NAD^+^ and ATP/ADP levels HD10.6 cells were treated with the indicated concentrations of either (A) piericidin A for six hours or (B) MGO for 24 hours. NADH and NAD were measured using the NAD/NADH-Glo assay. Boxes represent mean NADH/NAD^+^ and bars depict ± SEM. (n = 3 wells per group). n.s. = not significant, \**P* < 0.05, \*\**P* < 0.01, \*\*\*\**P* < 0.0001; One-way ANOVA and Dunnett’s multiple comparison test. (C) HD10.6 cells were treated with either vehicle, 500 nM piericidin A (6 hours), 1 µM MGO (24 hours), or 100 µM MGO (24 hours). To block glycolysis, 2-Deoxy-D-glucose (100 µM) was added in the last 6 hours of treatment. ATP and ADP were measured using the Sigma ATP/ADP ratio assay kit. Boxes represent mean ATP/ADP and bars depict ± SEM. (n = 3 wells per group). \**P* < 0.05, \*\**P* < 0.01; One-way ANOVA and Dunnett’s multiple comparison test.

**Fig. S3. Validation of *GCN2*^fl/fl^ *Pirt*^Cre/+^ line.** Left – Immunohistochemistry staining showing GCN2 expression in peripherin-positive cells. DRGs were harvested from either WT or *GCN2*^fl/fl^ *Pirt*^Cre/+^ animals and were stained for peripherin and GCN2. Right – Quantification of % GCN2 expression. Boxes represent mean % GCN2 expression and bars depict ± SEM. (n = 3 mice per group). \**P* < 0.05; Two-tailed unpaired t-test with Welch’s correction.

