## Supplement for "Mitochondrial dysfunction activates a translational stress response that drives diabetic pain"

Correspondence to: Zachary Campbell

**Short title:** GCN2 in diabetic pain

A

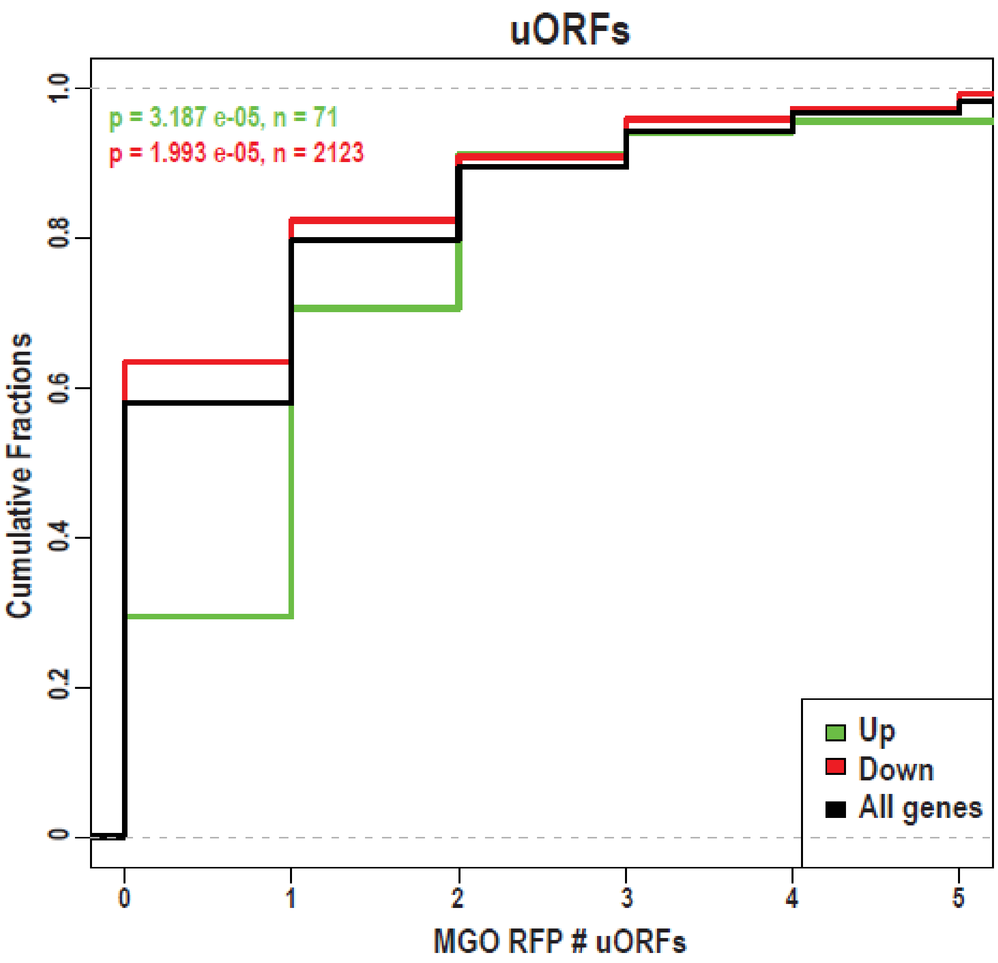


**Fig. S1. MGO upregulates the translation of uORF-containing transcripts**

Cumulative fraction plot showing the number of uORFs (MGO RFP # uORFs) across differentially expressed gene subsets compared to all evaluated genes. Genes upregulated upon MGO treatment (Up, green line; n = 71) show a significantly higher proportion of transcripts with multiple uORFs compared to background (All genes, black line). Genes downregulated upon treatment (Down, red line; n = 2,123) contain fewer uORFs relative to background. P values were obtained from Kolmogorov-Smirnov tests.

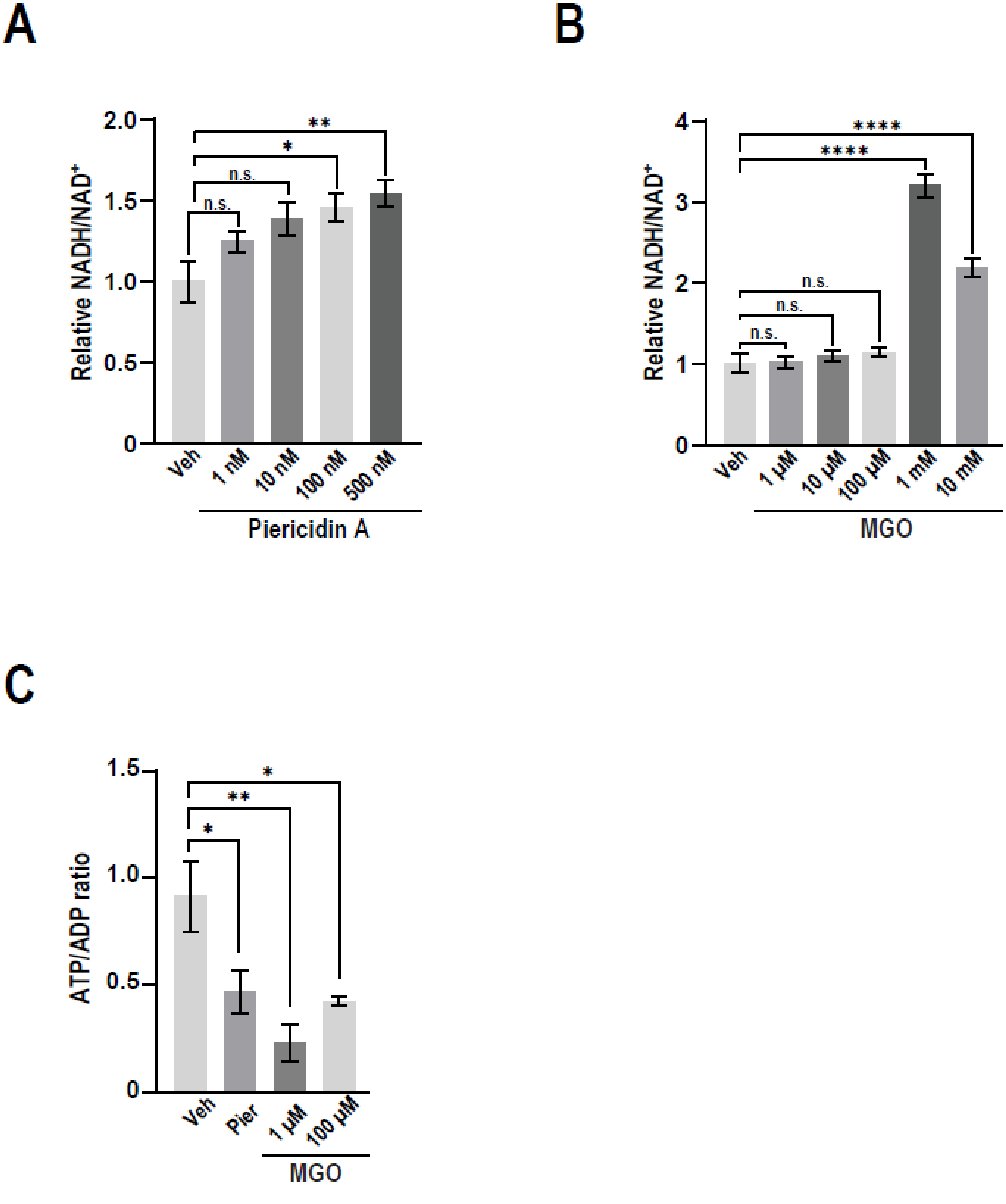


**Fig. S2. Piericidin A and MGO alter NAD/$NAD^+$ and ATP/ADP levels**

HD10.6 cells were treated with the indicated concentrations of either (A) piericidin A for six hours or (B) MGO for 24 hours. NADH and NAD were measured using the NAD/NADH-Glo assay. Boxes represent mean NADH/$NAD^+$ and bars depict ± SEM. (n = 3 wells per group). n.s. = not significant, *$P < 0.05$, **$P < 0.01$, ****$P < 0.0001$; One-way

ANOVA and Dunnett's multiple comparison test. (C) HD10.6 cells were treated with either vehicle, 500 nM piericidin A (6 hours), 1 µM MGO (24 hours), or 100 µM MGO (24 hours). To block glycolysis, 2-Deoxy-D-glucose (100 µM) was added in the last 6 hours of treatment. ATP and ADP were measured using the Sigma ATP/ADP ratio assay kit. Boxes represent mean ATP/ADP and bars depict ± SEM. (n = 3 wells per group). *$P$ < 0.05, **$P$ < 0.01; One-way ANOVA and Dunnett's multiple comparison test.

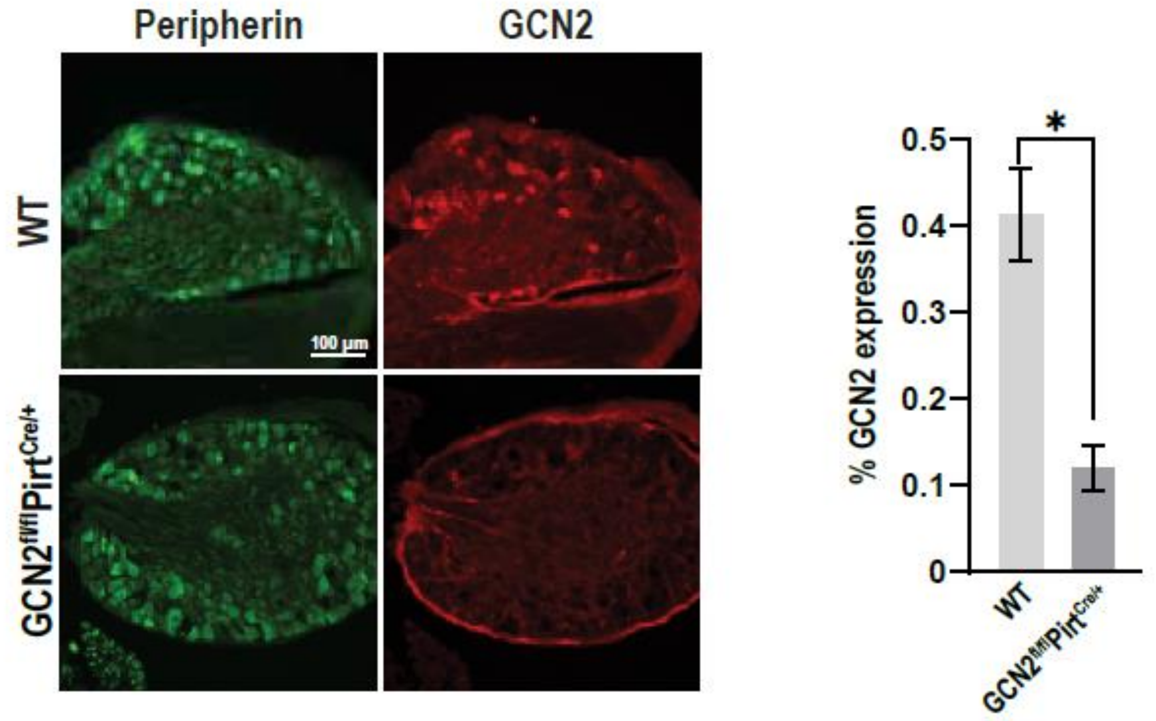


**Fig. S3. Validation of *GCN2*$^{fl/fl}$ *Pirt*$^{Cre/+}$ line.** Left – Immunohistochemistry staining showing GCN2 expression in peripherin-positive cells. DRGs were harvested from either WT or *GCN2*$^{fl/fl}$ *Pirt*$^{Cre/+}$ animals and were stained for peripherin and GCN2. Right – Quantification of % GCN2 expression. Boxes represent mean % GCN2 expression and bars depict ± SEM. (n = 3 mice per group). *$P < 0.05$; Two-tailed unpaired t-test with Welch's correction.
